# lncAPNet enables the deciphering of lncRNA–mRNA connections in patient transcriptomic data

**DOI:** 10.64898/2025.12.18.695074

**Authors:** Vasileios Vasileiou, George I. Gavriilidis, Pedro Faria Zeni, Marek Mraz, Karatzas Evangelos, Antonis Giakountis, Georgios A. Pavlopoulos, Antonis Giannakakis, Fotis Psomopoulos

**Affiliations:** Institute of Applied Biosciences, Center for Research and Technology Hellas, Thessaloniki, Greece; Department of Molecular Biology and Genetics, Democritus University of Thrace, Alexandroupolis, Greece; Berlin Institute of Health at Charité - Universitätsmedizin Berlin, Center of Digital Health, Berlin, Germany; Molecular Medicine, Central European Institute of Technology, Masaryk University, Brno, Czech Republic; European Molecular Biology Laboratory, European Bioinformatics Institute (EMBL-EBI), Wellcome Genome Campus, Hinxton, Cambridge, UK; Department of Biochemistry and Biotechnology, University of Thessaly, Larissa 41500, Greece; Institute for Fundamental Biomedical Research, BSRC “Alexander Fleming”, Vari 16672, Greece; Department of Computational Biology, Mohamed bin Zayed University of Artificial Intelligence (MBZUAI), Abu Dhabi, United Arab Emirates; University Research Institute of Maternal and Child Health and Precision Medicine, National and Kapodistrian University of Athens, 11527 Athens, Greece

## Abstract

**Motivation:** Long non-coding RNAs (lncRNAs) regulate gene expression through chromatin remodeling, transcriptional control, and post-transcriptional modulation, influencing physiological cell homeostasis but also disease onset. Yet most transcriptomic and network-based studies rely on descriptive linear co-expression analyses, missing nonlinear and mechanistic insights. Emerging ML/DL methods offer promise but remain limited by data sparsity, noise, insufficient biological priors, and poor interpretability, constraining systems-level lncRNA-mRNA motif discovery.

**Results:** In this manuscript, we introduce lncAPNet, an extended version of APNet workflow, which integrates graph-based nonlinear inference of lncRNA–mRNA interactions using NetBID2’s activity logic within an lncRNA-focused SJARACNe co-expression network, coupled with PASNet, a biologically informed sparse deep learning model. This framework enables explainable identification of lncRNA drivers in two different cancer type case studies, Chronic Lymphocytic Leukemia (CLL) and Prostate Adenocarcinoma (PRAD), uncovering lncRNA drivers that illuminate lncRNA-mediated programs in cancer progression.

**Availability and implementation:** lncAPNet’s R scripts, Python scripts, and methodologies are available at github repository: https://github.com/BiodataAnalysisGroup/lncAPNet

## 1. Introduction

Long non-coding RNAs (lncRNAs) are a diverse class of RNA molecules that are arbitrarily defined as longer than 200 nucleotides and play crucial roles in regulating gene expression and various cellular processes (Mattick et al. 2023). They function as signal drivers, decoys, guides, scaffolds, or enhancers to control transcriptional and post-transcriptional events (Zeni and Mraz 2021). Functionally, lncRNAs guide chromatin-modifying complexes to specific genomic loci for histone or DNA methylation, thereby influencing gene silencing or activation. They also regulate transcription by modulating RNA polymerase activity, post-transcriptional processes like splicing and mRNA stabilization, and translation by interacting with ribosomes or initiation factors. Moreover, some lncRNAs influence post-translational modifications by affecting protein acetylation or phosphorylation. Through these mechanisms, lncRNAs maintain genomic integrity, control cell differentiation, development, and signaling pathways. Dysregulation of lncRNA expression has been increasingly linked to the initiation and progression of various cancers, highlighting their potential as biomarkers and therapeutic targets in oncology (Nandwani, Rathore, and Datta 2021).

Transcriptomic studies of long non-coding RNAs (lncRNAs) primarily rely on bioinformatics approaches, with Differential Expression (DE) analysis and lncRNA–mRNA databases as the central strategy (Gong et al. 2021). However, they often remain descriptive, providing associations without directly uncovering the underlying molecular or regulatory functions of lncRNAs-mRNAs interactions or lncRNAs-pathways-related mechanisms.

Current lncRNA–mRNA network-level interactions approaches primarily rely on co-expression correlations when constructing lncRNA–mRNA networks using the Weighted Correlation Network Analysis (WGCNA) algorithm. Notably, many in silico studies suggest the importance of the correlation of lncRNAs with mRNA-pathway correlations (Xie, Shi, and Han 2023). In contrast, some computational studies (Hou et al. 2022) validated their computational results in vitro. However, due to the size of large-scale datasets and limited computational power, WGCNA cannot analyze all gene pairs and cannot capture nonlinear relationships between lncRNAs and mRNAs.

Despite their importance, the identification of lncRNAs in systems-level models in biology remains limited. The majority of ML/DL models in the domain, despite the ability to learn complex, non-linear, high-dimensional data patterns (Kourou et al. 2021), are not designed to capture the regulatory complexities or network-level behaviors of lncRNAs, nor do they incorporate biological priors essential for understanding their functional context (Kim and Lee 2023).

Many Deep Learning approaches are being tested to discover these nascent motifs behind lncRNAs and other omic modalities (Gong et al. 2021). Despite its potential, deep learning in lncRNA research faces major limitations, including data scarcity, high noise, and heterogeneity that compromise model robustness and reproducibility. Furthermore, incomplete and inconsistent information in existing lncRNA databases, together with the “black box” nature of deep learning models and the computational complexity of multiomics data integration, severely constrain interpretability, feature extraction, and biological validation (Kim and Lee 2023).

Considering the above, in the herein manuscript, we present lncAPNet, a novel computational pipeline built upon our recently reported Activity PASNet (APNet) framework (Gavriilidis et al. 2025). lncAPNet performs explainable AI (XAI)-driven supervised clustering of patient cohorts by integrating transcriptomic analysis with biological priors and the concept of differential activity. The original APNet integrates NetBID2 and SJARACNe workflows with a sparse, interpretable deep learning architecture (PASNet) for patient classification, supported by biological priors and post hoc visualization modules for clinical bioinformatics. In this extended version, lncAPNet expands the APNet toolbox by incorporating graph-based nonlinear interpretation of lncRNA–mRNA relationships through the SJARACNe/NetBID2 framework, embedding an lncRNA-specific knowledge graph (lncRNAlyzr-KG) into the biological priors (Evangelista et al. 2025), and unveiling lncRNA-associated regulons derived from the NetBID2 analysis. lncAPNet was tested in two distinct cancer pilot cases (Chronic Lymphocytic Leukemia – CLL and Prostate Adenocarcinoma – PRAD), in which our approach successfully captured both ground-truth and novel lncRNA-mRNA interactions, highlighting cancer hallmarks such as proliferation, tumor progression, metabolism, and malignancy.

## 2. Methodology

### 2.1 Concise overview of the modular architecture of lncAPNet

lncAPNet employs a three-stage pipeline. First, RNA-seq expression matrices are transformed into regulatory activity matrices using the SJARACNe/NetBID2 algorithm (Dong et al. 2023). Second, a biologically structured deep learning (DL) classifier based on the PASNet algorithm (Hao et al. 2018) is applied. Model explainability is achieved through SHAP (Shapley Additive Explanations) values (Aas, Jullum, and Løland 2021), which highlight the most informative lncRNAs, mRNAs, and their associated pathways. In the final stage, lncRNAs of interest are identified, and the lncHUB2 Appyters tool (Marino et al. 2023) is used to extract detailed information about these features, while Kaplan–Meier plots are generated to estimate the survival probability over time for each potential driver (Fig. 1).

**Figure 1:**
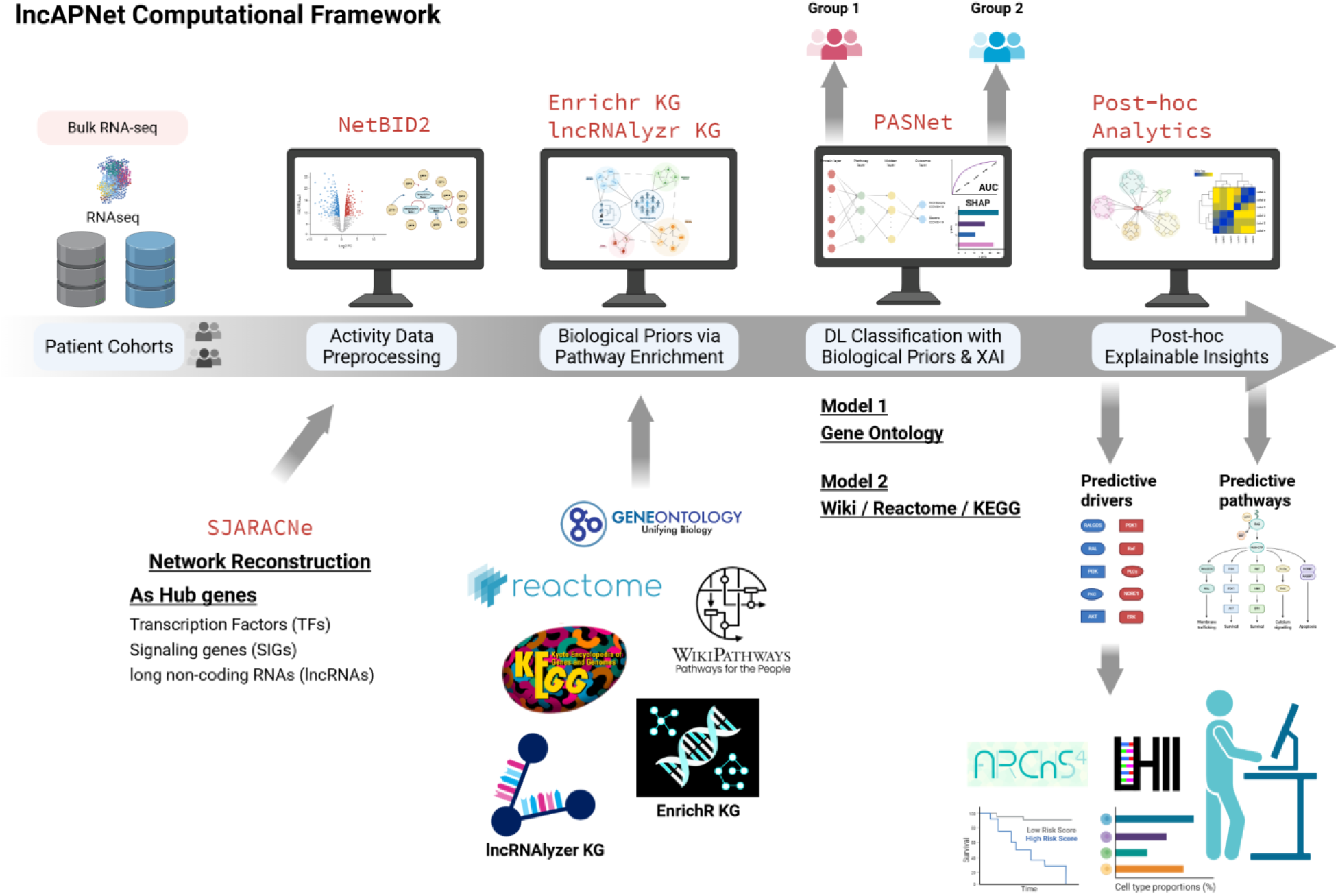
lncAPNet Overview. The lncAPNet workflow initially integrates NetBID2’s logic via the SJARACNe algorithm. Specifically, it identifies hub genes, including Transcription Factors (TFs), Signaling Genes (SIGs), and long non-coding RNAs (lncRNAs), and estimates their activity values using RNA-seq omics data. These activity values are then input into the PASNet algorithm, which leverages pathway information from GO, KEGG, Reactome, and WikiPathways as biological priors. Pathway–gene associations for mRNAs and lncRNAs are derived from the EnrichR-KG and lncRNAlyrs-KG databases developed by the Ma’ayan Lab, respectively. Finally, the critical features and pathways identified by PASNet are subjected to *post hoc* analysis using the open-source lncHUB2 tool to retrieve functional annotations for the relevant lncRNAs, while Kaplan–Meier plots are generated separately to identify potential biologically and prognostically meaningful drivers.

The key advancement of lncAPNet over the original APNet framework lies in its expanded initialization strategy: lncRNAs are designated as hub genes within the SJARACNe/NetBID2 algorithm, extending beyond the traditional focus on transcription factors (TFs) and signaling genes (SIGs). Additionally, lncAPNet incorporates pathway information linked to lncRNAs by leveraging the lncRNA-related knowledge graph developed by Ma’ayan’s Lab (lncRNAlyrs), thereby enriching the model’s biological context and interpretability. Finally, identified lncRNAs are validated using open-source annotated tools such as lncHUB2.

### 2.2 Activity values estimation and Driver inferences

For data formatting, we ensured a consistently formatted dataset by preprocessing the data. In this automated step, the inputs (count matrix, metadata, and optionally feature information) were converted into an ExpressionSet object consisting of a count matrix, sample metadata, and optionally feature annotations, which enabled using the NetBID2 pipeline for analyses.

The ExpressionSet objects of case studies used in this analysis were separately processed using the same workflow within the NetBID2 framework. Count matrix converted to log₂-formatted values to reduce data skewness and stabilize variance to prepare the data for subsequent statistical tests. Additional data preprocessing was also performed to discover the medium variability transcripts and informative samples for more detailed analysis.

**NetBID2** utilizes the **SJARACNe** algorithm to reverse-engineer context-specific interactomes, deriving “activity values” for each candidate driver gene. We define Transcription Factors (TFs), Signaling Genes (SIGs) from MSigDB, and long non-coding RNAs (lncRNAs) retrieved from BioMart as the pool of potential drivers for SJARACNe.

SJARACNe extends the original ARACNe method (Margolin et al. 2006) to handle large-scale data, assessing whether mutual information or correlation potentials are significantly different from zero (Khatamian et al. 2019). NetBID2 then leverages the SJARACNe output to calculate driver “activity values” as the weighted mean of each driver’s target genes:

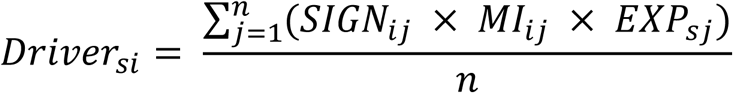

Where *EXPsj* is the expression values of target protein *j* in sample *s*, *MIij* is the mutual information between driver *i* and target *j*, and *SIGNij* is the sign of their Spearman correlation. Finally, *n* is the number of targets regulated by *i*.

Differential activity analysis is computed via the ‘getDE.BID.2G’ NetBID2 function.

### 2.3 Biological priors

For each case scenario, statistically significant up- and down-regulated (adj-pvalue < 0.05) drivers were mapped to biological pathways using the gseapy Python package, with gene–pathway associations derived from **EnrichR KG** for mRNAs (Evangelista et al. 2023) and **lncrnalyzr KG** for lncRNAs (Evangelista et al. 2025). The considered databases for the pathways enrichment use the Gene Ontology (GO), KEGG, Reactome, and Wikipathways. Pathways that pass the statistical significance cutoff were then depicted as one-hot encodings, where genes were assigned 0 if absent from a given pathway and 1 if present. Notably, because of the large number of GO pathways identified from the enrichment analysis, the modeling was carried out in two separate training runs: the first incorporating only the GO pathways, and the second including all remaining databases.

### 2.4 Deep Learning Neural-Network classification through PASNet-based biological priors

lncAPNet employs a sparse neural network, **PASNet** (Hao et al. 2018), for deep learning–based classification. PASNet integrates hierarchical driver-pathway relationships, allowing predictions to be both interpretable and explainable.

The mathematical foundation of PASNet involves enforcing layer-wise sparsity and incorporating cost-sensitive learning, which integrates hierarchical driver–pathway connections to produce more interpretable and explainable predictions. Specifically for the algorithmic background of PASNet:

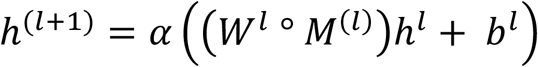

Where ○ is elementwise multiplication. For imbalanced data, the cost function includes class-specific average errors *Ck*. We optimize weights *W*(*l*) and biases *b*(*l*) via gradient updates:

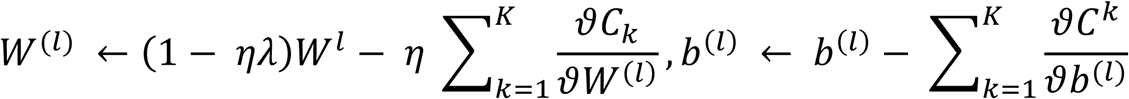

### 2.5 SHAP values

The **SHAP values** algorithm utilizes game theory to select the most interpretable feature drivers from the trained model. It computes a contribution score for each gene by quantifying its impact on the model prediction in every combination of drivers feasible, such that it gives a reasonable and consistent measure of importance. By adding up such contributions, drivers are ranked by their effect to gain insight into underlying biological processes and increase model interpretability. The mathematical rationale behind SHAP values is:

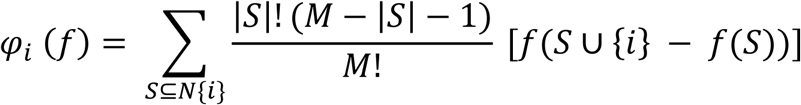

Where f is the input model and the N the number of input features.

### 2.6 Post Hoc explainability

Finally, lncAPNet offers diverse graph-based visualizations and analytical tools to extract meaningful biological insights. Feature importance is assessed using SHAP values, providing interpretable rankings of influential factors. Based on these rankings, the top predictive drivers can be selected to enhance the SJARACNe networks using driver-pathway correlations from PASNet, generating regulatory subgraphs that reveal key regulatory relationships. Specifically, lncAPNet can create driver–pathway bipartite graphs that combine multiple metrics (such as mutual information from SJARACNe-derived gene-gene correlations and PASNet learning gene-pathway weights), enabling researchers to filter and pinpoint subgraphs with potential translational significance.

To identify further information for the lncRNAs of interest, we deployed the lncHUB2 resources by using the lncHUB2 pipeline. Specifically, we utilized the lncHUB2 UMAP part to visualize lncRNA expression across 18,705 lncRNAs from ARCHS4 (Marino et al. 2023). As part of this process, the samples were first log2-transformed and quantile-normalized along the gene axis, after which UMAP was applied to the lncRNA expression data, treating samples as features. Moreover, the lncHUB2 resources provide a method to check whether the lncRNAs of interest are expressed in various tissues and cell types based on ARCHS4 data (Lachmann et al. 2018). Users can input specific lncRNAs along with tissues or cell types of interest to assess whether these lncRNAs show high expression through established tissues and cell types from databases. Finally, significant lncRNAs can be further analyzed using Kaplan-Meier plots (by using the lifelines Python package) to evaluate their association with patient survival, providing insights into their potential prognostic value.

APNet was developed following the **DOME guidelines** (Walsh et al. 2021) for reporting supervised machine learning analyses in biological studies (See Data and Code Availability).

## 3. Results

### 3.1 Case Studies

The two pilot studies differed in cancer type, design, and goals. The CLL study examined a hematologic cancer with heterogeneous outcomes, stratifying patients by IGHV status (Unmutated vs. Mutated) to capture prognostic differences. The PRAD study focused on a solid tumor and compared early (I–II) versus late (III–IV) clinical stages to assess progression. Both used lncAPNet, but the CLL study applied a cross-dataset PASNet approach with separate training and testing cohorts and incorporated survival data to identify prognostic lncRNAs. In contrast, the PRAD study evaluated a single cohort without survival analysis, identifying only stage-associated lncRNAs. Overall, the CLL pilot emphasized cross-cohort molecular drivers, while the PRAD pilot identified stage-related drivers within one dataset.

### 3.2 Case study 1: Chronic Lymphocytic Leukemia – CLL

#### CLL case study Overview

The first pilot case of lncAPNet pertains to CLL, one of the most frequent blood malignancies in the Western world. This leukemia is primarily dichotomised based on the mutational status of the IGHV gene in the B-cell receptor in indolent cases that require little to no treatment (Mutated-CLL; M-CLL) and in aggressive cases that can develop drug resistance with often fatal outcomes (Unmutated-CLL; U-CLL) (Fig. 2A) (Seda and Mraz 2015; Stevenson, Forconi, and Kipps 2021).

**Figure 2.**
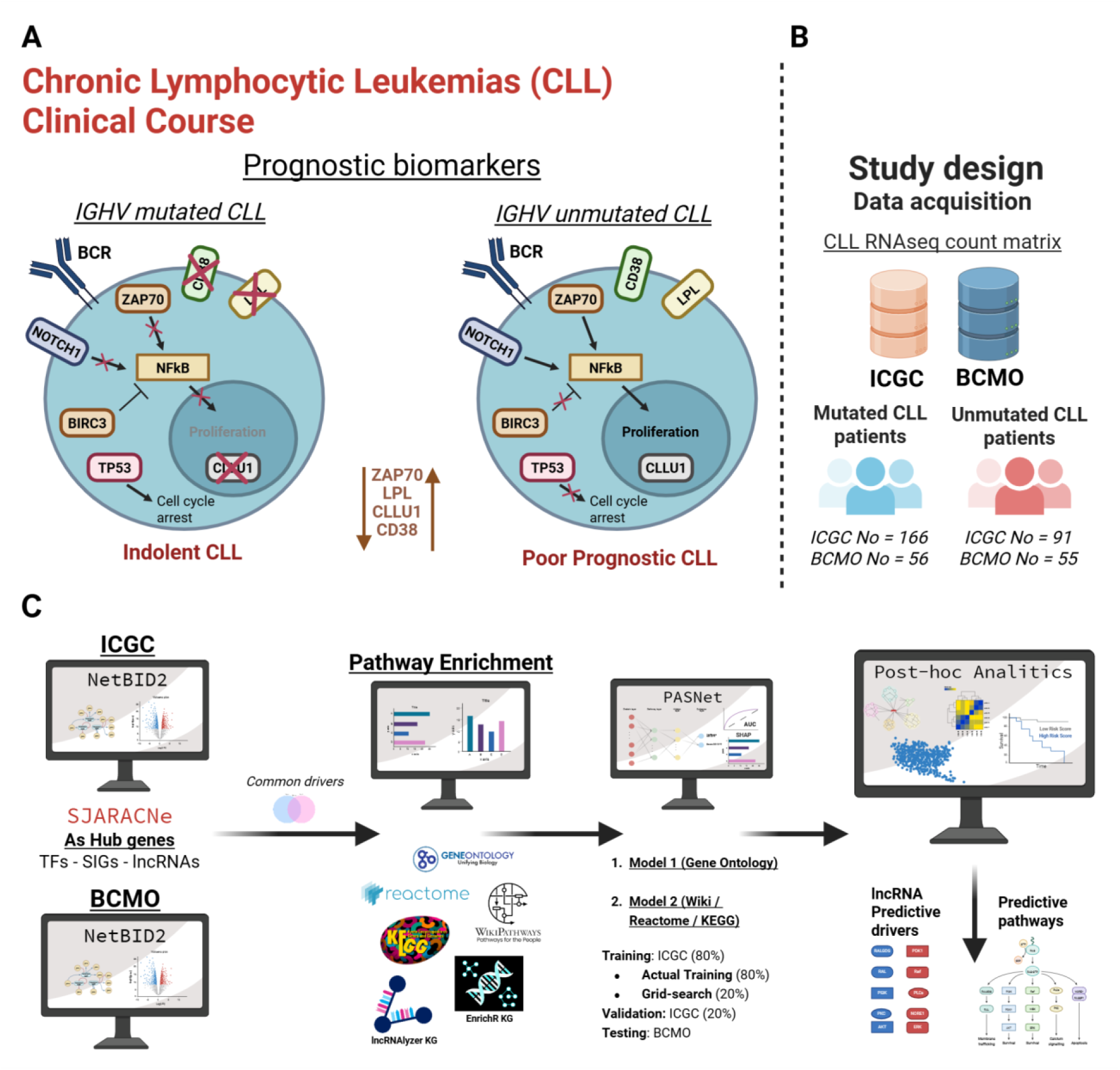
The lncAPNet framework applied to the CLL case study. (A) Brief explanation of the differences between Unmutated and Mutated CLL condition, including their known clinical prognostic biomarkers (based on (Rosenquist et al. 2013), and (B) a summary of the CLL patients in the dataset are stratified by IGHV status (U-CLL vs. M-CLL). (C) Οverview of lncAPNet applied in the CLL case-study scenario.

The primary objective of this analysis was to identify lncRNAs that are differentially active between U-CLL and M-CLL cases. To ensure robustness, two independent datasets, the International Cancer Genome Consortium (ICGC under the project CLLE-ES) and BloodCancerMultiOmics2017 Bioconductor database (BCMO) CLL cohorts, were initially analyzed separately using NetBID2, with one dataset designated as the training set and the other as the testing set within the PASNet framework. Both datasets included survival data, enabling downstream survival analyses of the lncRNAs identified.

#### Data retrieving and appropriate format creation

Bulk RNA-seq count matrices and their relevant clinical characteristics were first gathered from the ICGC and the BCMO (see Data Availability).

In the ICGC-CLL dataset, IGHV mutation status was manually extracted from associated metadata provided by the following study (Puente et al. 2015), whereas for the BCMO dataset, the mutation status was directly available within the relevant metadata (Dietrich et al. 2017).

The total number of samples for ICGC-CLL was 257 (split into 166 mutated and 91 unmutated IGHV status according to the study above) with a total amount of 38687 annotated transcript IDs, and the BCMO’s dataset had in total of 111 samples (split into 56 mutated and 55 unmutated IGHV status) with a total amount of 22915 annotated transcript IDs (Fig. 2B).

#### Activity Analysis and Identification of Driver Genes

We first applied the SJARACNe for the network reconstruction with their relevant gene lists (TFs, SIGs, lncRNAs), and ran separately for each dataset, by considering the default parameters of the SJARACNe algorithm (number of epochs at 100, consensus p-value threshold at 5×10⁻⁸).

Then, we used the NetBID2 algorithm to calculate gene activity and perform differential activity analysis independently on the ICGC and BCMO datasets. Differential analysis filtering was used to retain genes (adjusted p-value < 0.05 and target size > 30, suggested by the NetBID2 pipeline for robust drivers inference). Specifically, for the ICGC dataset, 1,921 drivers were identified, comprising 723 hyper-active and 1,198 hypo-active genes (793 overt and 1070 hidden drivers), whereas in the BCMO dataset, 3,551 significant drivers were identified, including 2,164 hyper-active and 1,387 hypo-active genes (1176 overt and 2261 hidden drivers). By categorizing drivers as positive (hyper-active) or negative (hypo-active) in each dataset, we identified 362 common positive and 212 common negative drivers, totalling 574 shared drivers (Sup. Fig. 1).

#### Pathway Enrichment Analysis

To investigate the functional relevance of these drivers, pathway enrichment analysis was performed on the common hyperactive and hypoactive drivers using GO, Reactome, KEGG, and Wikipathways. Pathways were considered significant at an adjusted p-value < 0.05, resulting in the identification of 256, 103, 45, and 28 pathways, respectively.

Given the high number of GO pathways, we conducted two separate analyses/model training: (i) GO enrichment with 256 pathways, involving 229 of 574 drivers, and (ii) enrichment across KEGG, Reactome, and Wikipathways with 176 pathways, involving 269 of 574 drivers (Sup. Fig. 2).

#### Model Training and Validation

The ICGC dataset was partitioned into three subsets: first, an 80–20% split was applied to generate the training set and the validation set. The training part (80%) was then further divided into training and grid-search validation sets in an 80–20% ratio. For grinding, the second training dataset was used, and the grinding part was validated for the optimal Learning rate (LR) and L2 regularization optimal parameters at 10.000 epochs, whereas those parameters were used for the actual training by using the training part and the second validation part of the dataset. After model training, performance was evaluated on an independent test dataset, BCMO. Those steps were followed by both model training.

For GO-based analysis, the ICGC-training and ICGC-grinding validation datasets yielded an optimal learning rate (LR = 0.005) and L2 regularization (L2 = 0.0005). Subsequently, lncAPNet was trained and validated on the ICGC datasets, achieving robust performance metrics (AUC = 0.96, F1 score = 0.86). Applying this model to the independent BCMO dataset produced similarly high metrics (AUC = 0.96, F1 score = 0.84) (Fig. 3A).

**Figure 3.**
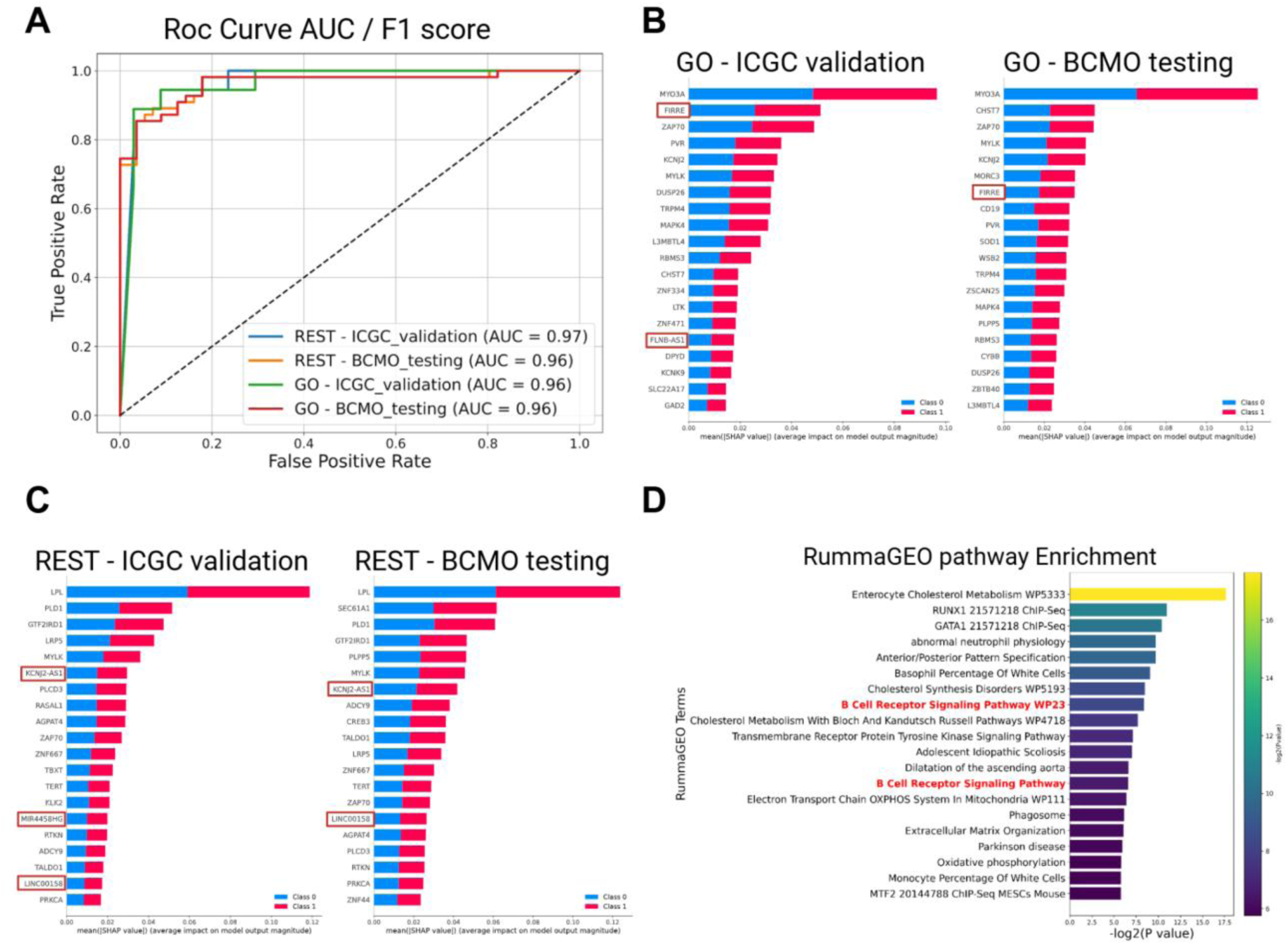
lncAPNet results in the CLL case study. (A) ROC curves with AUC and F1 scores for different lncAPNet configurations, including ICGC validation using the GO database, ICGC validation using additional databases, BCMO testing with the GO database, and BCMO testing with the remaining databases. (B-C) Top 20 SHAP features displayed as bar plots for each comparison scenario: ICGC validation database and BCMO testing with the GO database (B), ICGC validation with other databases, and BCMO testing with the rest databases (C). (D) Pathway enrichment analysis (RummaGEO) based on top SHAP features, with terms associated with both CLL and IGHV status highlighted in red on the y-axis.

For the combined KEGG, Reactome, and Wikipathways analysis, optimal hyperparameters were LR = 0.007 and L2 = 0.0005. Training and validation on ICGC datasets achieved AUC = 0.97 and F1 score = 0.86, with external validation on the BCMO dataset yielding AUC = 0.96 and F1 score = 0.84 (Fig. 3A).

#### Identification of Key lncRNA Drivers

For each model, a SHAP analysis was conducted to identify the top 20 features that contribute to the model’s predictions (Fig. 3B-C). By combining the SHAP values from both models, 49 unique significant explainable drivers were identified, including five lncRNAs: FIRRE, MIR4458HG, LINC00158, KCNJ2-AS1, and FLNB-AS1. Enrichment analysis using RummaGEO demonstrated that those top 49 drivers were involved in BCR signaling (Fig. 3D).

#### Bipartite Graphs driver-pathway correlations

After extracting the driver-pathway correlations from PASNet, we exclude those weights within the interquartile range (25th–75th percentile) to focus on the most relevant correlations. We then integrated the outputs from both GO and other pathway databases (KEGG, Reactome, and WikiPathways), and applied a filtering criterion, retaining pathways associated with at least three drivers and drivers participating in at least one pathway. Through this approach, we identified four lncRNAs [FIRRE, LINC00158, KCNJ2-AS1, and FLNB-AS1] that exhibited significant correlations with these pathways (Sup. Fig. 3).

The graph is organized into five modules, each reflecting a major biological process. Module 1 centers on transcriptional regulation, within which FIRRE appears as a subgraph. Module 2 represents a MAPK–RAS–RTK signaling hub that governs proliferation, migration, and growth-factor responses, and includes FLNB-AS1 as a subgraph. Module 3 involves protein phosphorylation and receptor tyrosine kinase signaling, supporting cellular responses to external stimuli, with KCNJ2-AS1 embedded as a subgraph. Module 4 combines immune and metabolic pathways—such as cytokine signaling and carbohydrate metabolism—and contains LINC00158 as a subgraph. Finally, Module 5 integrates neuronal, hormonal, and gene-expression pathways, including glutamatergic synapse and GnRH signaling (Fig. 4A).

**Figure 4.**
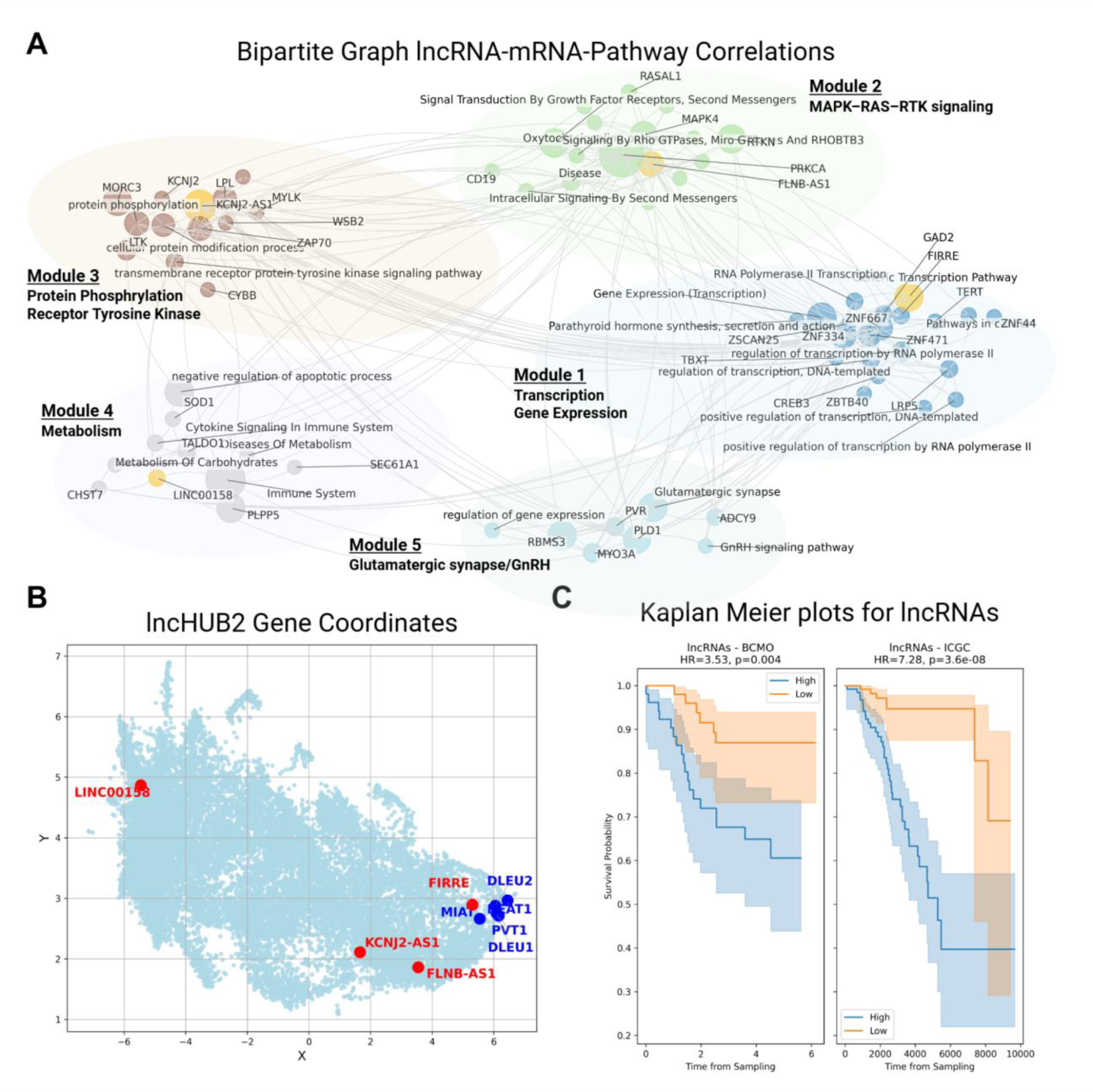
Post-hoc analysis applied to the CLL case study. (A) Bipartite graph showing correlations among drivers and between drivers and pathways, and clustered by the community-leiden algorithm. lncRNAs were colored with yellow-orange. (C) UMAP visualization from lncHUB2 showing how close the lncRNAs identified in the current analysis (red) with those lncRNAs previously reported in CLL studies (blue). (D) Kaplan–Meier survival plots for each dataset (ICGC and BCMO) using lncRNAs selected from the top SHAP values, incorporating survival status and lncRNA activity values.

#### Validation with ground-truth lncRNAs in CLL

Using the lncHUB2 pipeline and examining the UMAP projection, we observed that several lncRNAs from our analysis cluster closely with previously reported CLL-associated lncRNAs (Fig. 4B). In particular, FIRRE were found to be statistically closer to the known CLL-related lncRNAs, with observed average distances of 0.332 (Sup. Table 1).

In contrast, other lncRNAs from our dataset showed associations with broader cell types and tissues. This suggests that although not all lncRNAs are closely associated with the established CLL-associated cluster, those enriched in lymphoid tissues and lymphocyte-related cell types may still play biologically significant roles in CLL pathogenesis and warrant further investigation. Specifically, FIRRE and LINC00158 exhibited high expression in lymphoblastic and lymphoblastoid tissues. Moreover, FIRRE, LINC00158, and KCNJ2-

AS1 showed elevated expression in hematopoietic and lymphoid-derived cell lines, including REH, HL60, Raji, and BJAB. These patterns highlight their potential relevance in lymphoid biology and suggest functional involvement in CLL-related cellular contexts (Sup. Fig. 4).

#### Prognostic value of discovered lncRNAs in CLL

Given the most significant lncRNA drivers identified in the previous analysis, the samples were split into equal groups (50%-50%) based on whether each lncRNA was higher or lower in activity, and their survival was evaluated using Kaplan-Meier plots. Survival status was retained, and p-values and hazard ratios (HR) were calculated for each dataset based on activity values, following the NetBID2 approach for Kaplan–Meier plots using activity values. Notably, all selected lncRNAs appeared to be critical across both datasets (Sup. Fig. 5). Moreover, when all lncRNAs were combined to assess their collective contribution to patient survival, the Kaplan–Meier analysis revealed significant associations in both datasets: BCMO (HR = 3.53, p = 0.004) and ICGC (HR = 7.69, p = 1.1 × 10⁻⁸) (Fig. 4C).

### 3.3 Case study 2: Prostate Adenocarcinoma – PRAD

#### PRAD case study Overview

The second scenario focused on PRAD, in which patients were stratified according to tumor stage, distinguishing early-stage (stage I–II) from late-stage (stage III–IV) disease to reflect increasing tumor burden, aggressiveness, and metastatic potential (Fig. 5A). Clinical characteristics were estimated for prostate cancer, including the Gleason score, which is derived as the sum of the individual Gleason grades (primary and secondary patterns). The N and M classifications serve as more specific indicators of disease progression, denoting lymphatic involvement and distant metastasis, respectively. In combination, these variables contribute to the determination of the overall disease stage (Cheng et al. 2012). The objective was to identify lncRNAs associated with tumor progression and malignancy, capturing molecular changes linked to advancing disease. Although comprehensive clinical data for survival analyses were not available in this cohort, lncAPNet was successfully applied to extract candidate lncRNA drivers correlated with cancer stage, providing insights for tumor progression.

**Figure 5.**
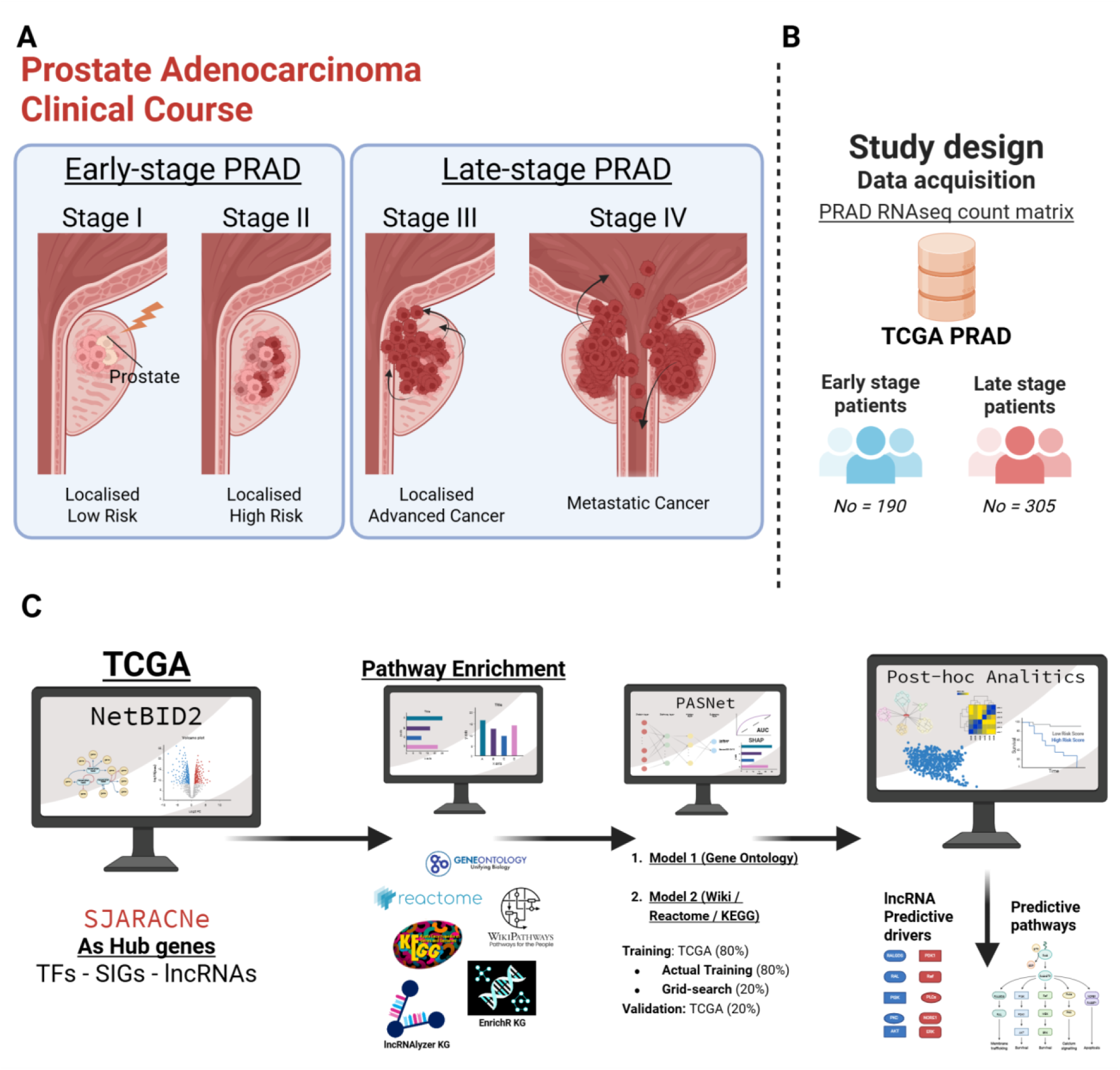
The lncAPNet framework applied to the PRAD case study. (A) A concise overview of PRAD progression, summarizing the key pathological and clinical features that characterize each stage of the disease. (B) Summarize the patients that stratified into Early and Late stages according to the metadata. (C) A schematic overview of the lncAPNet framework is applied in the PRAD case-study scenario.

#### Data retrieving and appropriate format creation

PRAD bulk RNAseq count matrix and metadata were retrieved from The Cancer Genome Atlas (TCGA) via the Genomic Data Commons (GDC) Data Portal, through the TCGA-PRAD project. The clinical characteristics of the samples were categorized into early (Stage I, II) and late (Stage III, IV) stages of cancer, indicating the malignancy of the cancer. Specifically, the number of early-stage cancer patients was *Ne = 190*, whereas the number of late-stage patients was *Nl = 305*. Finally, after filtering and transcript annotation, the total number of annotated transcript IDs was 33403 (Fig. 5B).

#### Activity Analysis and Identification of Driver Genes

The second case study focuses on the malignancy of PRAD, splitting the dataset into early stages of cancer (Stage I, II) and late stages of cancer (Stage III, IV).

Similarly to the first case study scenario, we applied the SJARACNe algorithm for network reconstruction using its relevant gene lists (TFs, SIGs, lncRNAs), with the default parameters of the algorithm (number of epochs set to 100, consensus p-value threshold at 5 × 10⁻⁸).

The NetBID2 algorithm was applied again to the entire RNA-seq dataset, comparing the two groups as described above. Similarly to the previous case study scenario, we kept the drivers that have adj-pvalue < 0.05 and at least 30 target genes (as recommended by the NetBID2 pipeline for reliable drivers inference) from the SJARACNe algorithm. Notably, the significant drivers that were extracted from this filtering were 4331 [2006 hyperactive and 2325 hypoactive drivers]. Interestingly, only 2.609 of those drivers were noticed as overt drivers, those that are important with the differential Expression method, whereas 1.722 of the mentioned drivers were counted as hidden, whose expression analysis was not statistically significant (Sup. Fig. 6).

#### Pathway Enrichment Analysis

In a total of 4331 significant drivers, we applied Pathway Enrichment analysis and set a cutoff of adjusted p-values at 0.05. We extracted 140, 14, 92, and 0 pathways from GO, KEGG, Reactome, and Wikipathways, respectively.

Due to the large number of GO pathways, the analysis was separated into two distinct model training scenarios: (i) GO with 140 pathways, involving 1756 out of 4331 drivers, and (ii) with the rest of the databases, with 106 pathways, involving 1490 out of 4331 drivers (Sup. Fig. 7).

#### Model Training and Validation

Similarly, for the CLL scenario, the dataset was first split 80–20% into a training set and a validation set. The 80% training portion was then further divided into training and grid-search validation subsets using an 80–20% split. For the grid search, the second training subset was used, and the grid-search validation subset was employed to determine the optimal Learning rate (LR) and L2 regularization parameters at 10.000 epochs. These optimized parameters were then applied to the final model training using the refined training and validation subsets.

For GO-based analysis, the TCGA-training and TCGA-grinding validation datasets yielded an optimal learning rate (LR = 0.01) and L2 regularization (L2 = 0.0003). Subsequently, lncAPNet was trained and validated on the TCGA datasets, achieving robust performance metrics (AUC = 0.84, F1 score = 0.78). (Fig. 6A).

**Figure 6.**
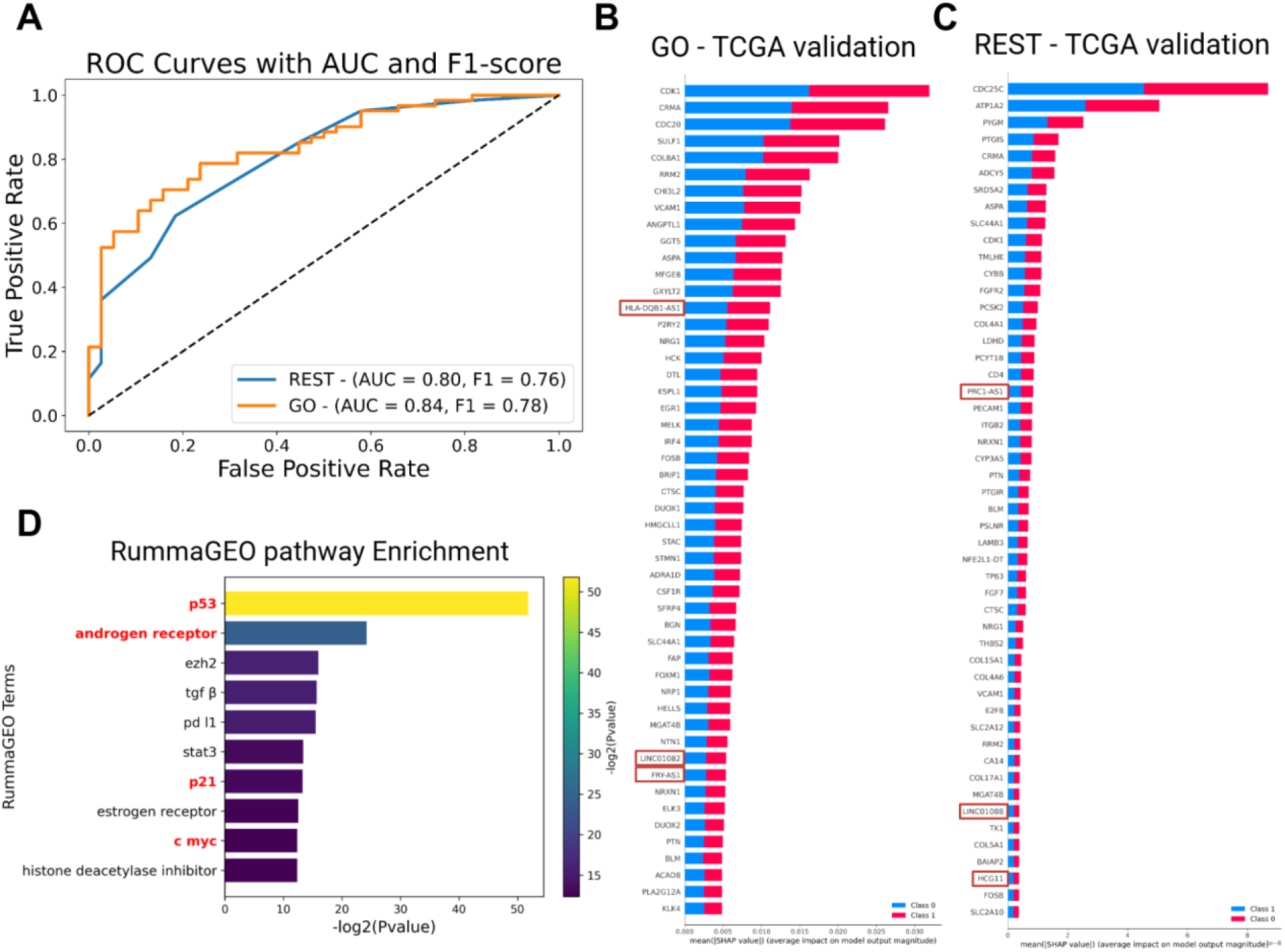
lncAPNet results in the PRAD case study. (A) ROC curves showing AUC and F1 scores for different lncAPNet configurations, including validation using the GO database and validation using other databases. (B-C) Top 50 SHAP features displayed as bar plots for each comparison scenario: TCGA validation with the GO database (B) and TCGA validation with the remaining databases (C) (KEGG, Reactome, and WikiPathways). (D) Pathway enrichment analysis (RummaGEO) based on the top SHAP features from all analyses, with terms associated with PRAD highlighted in red on the y-axis.

For the combined KEGG, Reactome, and Wikipathways analysis, optimal hyperparameters were LR = 0.007 and L2 = 0.0003, whereas the validation on TCGA datasets achieved AUC = 0.79 and F1 score = 0.76 (Fig. 6A).

#### Identification of Key lncRNA Drivers

In this step, SHAP was employed to identify the lncRNA features most influential in model predictions. For each model, the top 50 features ranked by SHAP values were extracted and combined into a unique list of significant features (87 drivers) (Fig. 6B-C). Notably, six lncRNAs were consistently ranked among the highest-scoring features: PRC1-AS1, FRY-AS1, LINC0182, HLA-DQB1-AS1, LINC01088, and HCG11. Enrichment analysis using RummaGEO indicated that these top-ranked drivers were linked to biological molecules implicated in the disease (Fig. 6D).

#### Bipartite Graphs driver-pathway correlations

Following the extraction of driver-pathway correlations from PASNet, we focused on the most meaningful associations by excluding those within the interquartile range (25th–75th percentile). Next, we combined results from GO with additional pathway databases, including KEGG, Reactome, and WikiPathways, and implemented a filtering step: retaining only pathways linked to at least three drivers and drivers involved in at least one pathway. Using this strategy, we pinpointed four lncRNAs [PRC1-AS1, FRY-AS1, LINC0182, and HCG11] that showed strong correlations with these pathways (Sup. Fig. 8).

The network organizes into distinct clusters that represent key biological processes. Module 1, which includes FRY-AS1, LINC0182, and HCG11, is enriched for transcriptional regulation and RTK signaling, featuring pathways such as RNA Polymerase II transcription, regulation of transcription, and signaling by receptor tyrosine kinases. Module 2 centers on metabolic processes, encompassing glycerophospholipid biosynthesis, phospholipid metabolism, amino acid catabolism, and pathways related to the extracellular matrix and vascular transport. Module 3 highlights extracellular matrix and collagen organization, including collagen biosynthesis and modifying enzymes, collagen chain trimerization, and extracellular structure organization. Module 4, which contains PRC1-AS1, focuses on the cell cycle and phosphorylation, with pathways such as G1/S-specific transcription, gene expression, and protein phosphorylation. Finally, Module 5 involves immune signaling and protein modification (Fig. 7A).

**Figure 7.**
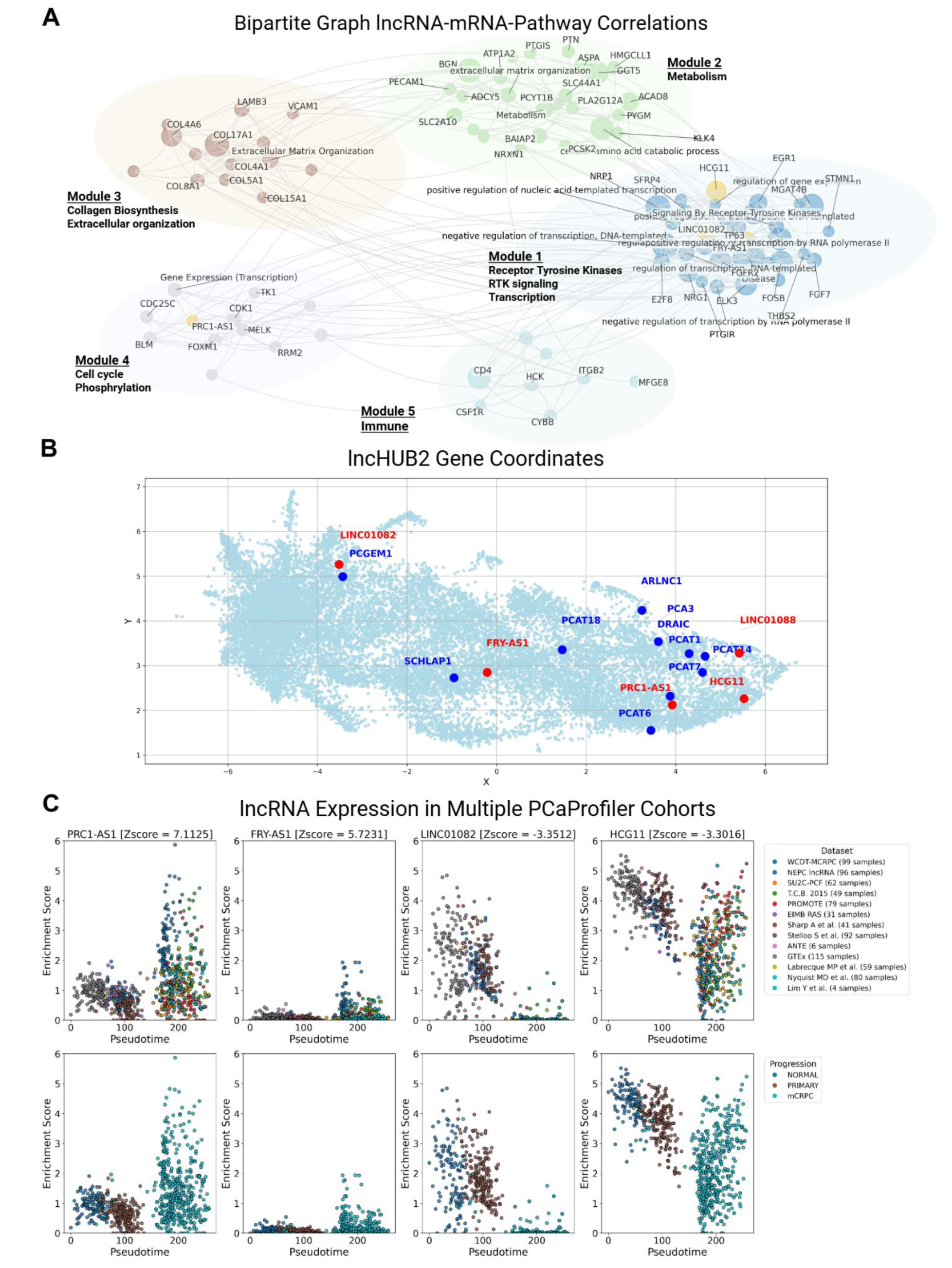
Post-hoc analysis applied to the PRAD case study. (A) A bipartite graph illustrates the correlations among driver genes as well as between drivers and pathways, with communities identified using the Leiden clustering algorithm; lncRNAs are highlighted in yellow–orange for visual distinction. (B) A UMAP projection from lncHUB2 displays the spatial relationship between lncRNAs identified in the current analysis (red) and lncRNAs previously reported in prostate cancer studies (blue), showing their relative proximity in expression space. (C) Scatter plots derived from the PCaProfiler demonstrate, using independent datasets (upper panels), how the expression of the lncRNAs of interest changes along prostate cancer progression inferred through pseudotime analysis (lower panels). For each lncRNA, its corresponding Z-score from the differential activity analysis is also provided.

#### Validation with ground-truth lncRNAs in PRAD

Using the lncHUB2 pipeline and examining the UMAP projection, we observed that several lncRNAs from our analysis cluster closely with previously reported Prostate cancer-associated lncRNAs (Fig. 7B). In particular, PRC1-AS1 and LINC01082 were found to be statistically closer to the known PRAD-related lncRNAs, with observed average distances of 0.200 and 0.284, respectively (Sup. Table 2).

In contrast, other lncRNAs in our dataset exhibited associations with a broader range of cell types and tissues. This indicates that, although not all lncRNAs are closely linked to the established prostate-associated cluster, PRC1-AS1 and LINC01082 are enriched in sperm and spermatozoa, whereas HCG11 shows enrichment in the urogenital reproductive system. Additionally, HCG11 correlates with prostate-related cell lines such as DU-145 and FRY-AS1 with C4-2. LINC01082 is also detectable in PC-3 cells, though its expression is very low and likely negligible (Sup. Fig. 9).

Pseudotime in the Prostate Cancer Atlas (PCaProfiler) represents an inferred molecular trajectory that orders samples from normal to advanced disease based on global gene expression patterns. It captures gradual transcriptional changes that occur during prostate cancer progression rather than reflecting actual chronological time. Enrichment scores, calculated using ssGSEA, quantify pathway or gene-signature activity within each sample. Along pseudotime, PRC1-AS1 and FRY-AS1 show increasing enrichment, indicating elevated activity in advanced stages, whereas HCG11 decreases and LINC01082 becomes nearly absent. These patterns align with differential activity Z-scores, where PRC1-AS1 and FRY-AS1 are positively active, and HCG11 and LINC01082 show negative activity. Together, these findings highlight stage-specific lncRNA dynamics and their potential roles in promoting prostate cancer progression (Fig. 7C).

## 4. Discussion

The therapeutic potential of lncRNAs has garnered significant attention in recent years, with several candidates advancing toward clinical evaluation. lncRNA expression can be modulated through multiple approaches, including siRNAs, antisense oligonucleotides, CRISPR-Cas9, and small molecule inhibitors (Tamblin-Hopper et al. 2024), with several preclinical programs demonstrating promising results. Notably, HAYA Therapeutics is advancing HTX-001, which targets the lncRNA WISPER, a master regulator of cardiac fibrosis, toward clinical trials for heart failure treatment. The tissue and cell-type specificity of lncRNAs makes them particularly attractive therapeutic targets, as their restricted spatial and temporal expression patterns enable lineage-specific gene therapy with potentially reduced off-target effects (Tamblin-Hopper et al. 2024). Well-characterized lncRNAs such as MALAT1 (Puri, Majumder, and Gaikwad 2025), HOTAIR (Zhou et al. 2015), and ANRIL (Yin et al. 2021), which have been implicated in cancer progression, cardiovascular disease, and inflammatory conditions, exemplify the breadth of diseases amenable to lncRNA-targeted interventions.

Our pipeline’s ability to systematically identify lncRNA-mRNA-pathway interactions provides a framework for prioritizing candidate lncRNAs based on their regulatory roles in disease-relevant pathways, potentially accelerating the discovery of novel therapeutic targets. While lncRNAlyzr represents a valuable resource for exploring lncRNA-mRNA relationships through its knowledge graph structure, its current implementation remains relatively static, limiting its utility for data-driven biological discovery and clinical decision-making. To address this limitation, we propose integrating deep learning approaches that can leverage the knowledge graph architecture while maintaining biological interpretability. By enabling deep learning models to access and learn from the knowledge graph’s structured relationships, we can facilitate biologically informed decisions such as supervised patient stratification, where the model not only performs clinical classifications but also reveals which specific lncRNA-mRNA connections drive its predictions. This integration forms the basis of lncAPNet, a comprehensive framework that synergistically combines knowledge graph relational connections, co-expression networks derived from SJARACNe, deep learning classification models, explainable AI through SHAP (SHapley Additive exPlanations) values, and network analytics. This multi-layered approach enables researchers to move beyond simple database queries toward a mechanistic understanding, identifying the key lncRNA-mRNA regulatory axes that underlie disease phenotypes and potentially guiding the selection of lncRNA therapeutic targets based on their functional relevance in patient subgroups.

The cornerstone of lncAPNet lies in the activity inference strategy that relies on both the activity values and the biological priors (knowledge), specifically focusing on lncRNAs. As emphasized in the original APNet (Gavriilidis et al. 2025), activity values provide two key advantages: (i) better reflect regulatory dependencies between lncRNAs and protein-coding molecular entities, and (ii) enable the extraction of biologically meaningful lncRNA driver–pathway relationships. These properties are particularly crucial for lncRNAs, whose functions are often mediated through complex, nonlinear interactions with mRNAs.

In the CLL case study, the analysis identified a set of significantly dysregulated signaling and survival pathways correlated with the unmutated IGHV phenotype. These include heightened activation of the B-cell receptor (BCR) (Chen et al. 2024; 2008) and downstream tyrosine kinase–PI3K/AKT/MAPK/Ras pathways (Murali et al. 2021), enhanced cytokine and immune regulatory signaling (e.g., IL-4, CD40) (Coscia et al. 2010), increased engagement of the anti-apoptotic BCL2/NF-κB survival program (Majid et al. 2008), and greater responsiveness to microenvironmental and migration cues. Notably, genes such as LPL, ZAP70, PRKCA, and CD19 (Álvarez-Silva et al. 2015; Chen et al. 2024; Maloum et al. 2009) were identified as critical molecules of disease progression in the unmutated IGHV status.

Among the significant lncRNAs, FIRRE emerged as a potential key driver across the analysis, demonstrating a strong association with both survival status and CLL-related lncRNA networks. According to existing literature, FIRRE, through the interaction with PTBP1, which is activating the Wnt/β-catenin pathway, contributes to stabilizing BCR signaling in DLBCL, thereby promoting B cell proliferation and malignant progression (Liu et al. 2022).

In late-stage PRAD case studies, results show that correlated pathways and genes, including dysregulated receptor tyrosine kinase signaling driven by FGFR2, FGF7, and NRP1, activate PI3K/AKT (Wang et al. 2024) and MAPK pathways (Shen et al. 2021) to sustain proliferation, while CDK1, CDC25C, and TK1 promote cell cycle progression and genomic stability (Xie et al. 2022). ECM remodeling and collagen biosynthesis (Stewart, Cooper, and Sikes 2004), mediated by COL4A1, COL5A1, COL8A1, COL15A1, and THBS2, enhance invasion and angiogenesis, complemented by transcriptional regulators FOXM1 (Liu et al. 2017), MELK/E2F8 (Lee et al. 2023), and FOSB (Barrett, Millena, and Khan 2017), which drive oncogenic programs. Secreted and adhesion molecules such as KLK4 (Klokk et al. 2007), PTN (Liu et al. 2021), VCAM1 (Chen et al. 2015), and CSF1R (Escamilla et al. 2015) further support angiogenesis and immune evasion, collectively promoting metastasis and tumor aggressiveness.

Notably, the lncRNAs PRC1-AS1, HCG11, LINC01082, and FRY-AS1 were identified as significant findings, potentially acting as key regulatory molecules within these oncogenic networks. Notably, reduced expression of the lncRNA HCG11 has been associated with the promotion of prostate cancer progression (Zhang et al. 2016) and has also been reported to inhibit the phosphoinositide 3-kinase/protein kinase B (PI3K/AKT) signaling pathway, thereby suppressing prostate cancer progression (Wang et al. 2019). Moreover, LINC01082 has been proposed as a potential prognostic biomarker, as it appears to play a critical role in patient survival (Zhang et al. 2022). Consistently, downregulation of LINC01082 promoted 22RV1 cell migration, suggesting that LINC01082 may suppress prostate cancer cell motility by upregulating or activating DUSP2. These findings indicate that LINC01082 exerts its tumor-suppressive effects, at least in part, through the DUSP2-mediated pathway, potentially linked to the FOXA1-associated ceRNA network (Yang et al. 2023).

## 5. Conclusions and Future Work

Despite its advantages, lncAPNet has several limitations in explaining biological mechanisms. First, the reliability of prior knowledge depends heavily on the comprehensiveness of current lncRNA databases, which remain less mature than those available for protein-coding genes. Second, functional interpretation of predicted lncRNA drivers must be approached with caution, as many lncRNAs are highly tissue- and context-specific. Third, co-expression does not necessarily imply direct interactions with their protein-coding targets; rather, it provides a broader view of the pathways in which they participate. Finally, the current script-based implementation may limit scalability in cloud or HPC environments, where workflow managers such as Nextflow could facilitate deployment.

Looking forward, lncAPNet can be generalized across multiple cancer-disease settings, particularly where lncRNAs have already been implicated as biomarkers or therapeutic targets. Its activity-based integration of lncRNAs into DL frameworks offers a modular and expandable strategy: future iterations could incorporate single-cell and/or spatial multi-omics to capture the cellular and spatial heterogeneity of lncRNA regulation. Furthermore, in vitro and in vivo validation of predicted lncRNA drivers will be essential to establish their translational value.

Overall, lncAPNet represents a conceptual and technical advance in interpretable deep learning for systems biology. By positioning lncRNAs as hub regulators within the activity-driven framework of APNet, it bridges the coding and non-coding layers of regulation, thereby improving both predictive power and mechanistic insight.

## Data and Code Availability

The lncAPNet R and Python scripts of the workflow of this manuscript can be found at: https://github.com/BiodataAnalysisGroup/lncAPNet

The RNAseq count matrices and their relevant metadata that were used in this study can be accessed in the following links

- ICGC ARGO [CLLE-ES]: https://platform.icgc-argo.org/
- BloodCancerMultiOmics2017 R package: https://bioconductor.org/packages/release/data/experiment/html/BloodCancerMultiOmics2017.html
- TCGA PRAD: https://portal.gdc.cancer.gov/projects/TCGA-PRAD All ExpressionSet formats of the above data are also included in our Zenodo: https://doi.org/10.5281/zenodo.17786260

DOME Registry: https://identifiers.org/dome:xpjtohmv78

## Abbreviations

lncRNAs: Long non-coding RNAs
DE: Differential Expression
DA: Differential Activity
BCMO: BloodCancerMultiOmics2017
ML: Machine Learning
DL: Deep Learning
GO: Gene Ontology
CLL: Chronic Lymphocytic Leukemia
U-CLL: Unmutated CLL
M-CLL: Mutated CLL
PRAD: Prostate Adenocarcinoma
PCaProfiler: Prostate Cancer Atlas

## Supplementary Materials

**Supplementary Figure 1.**
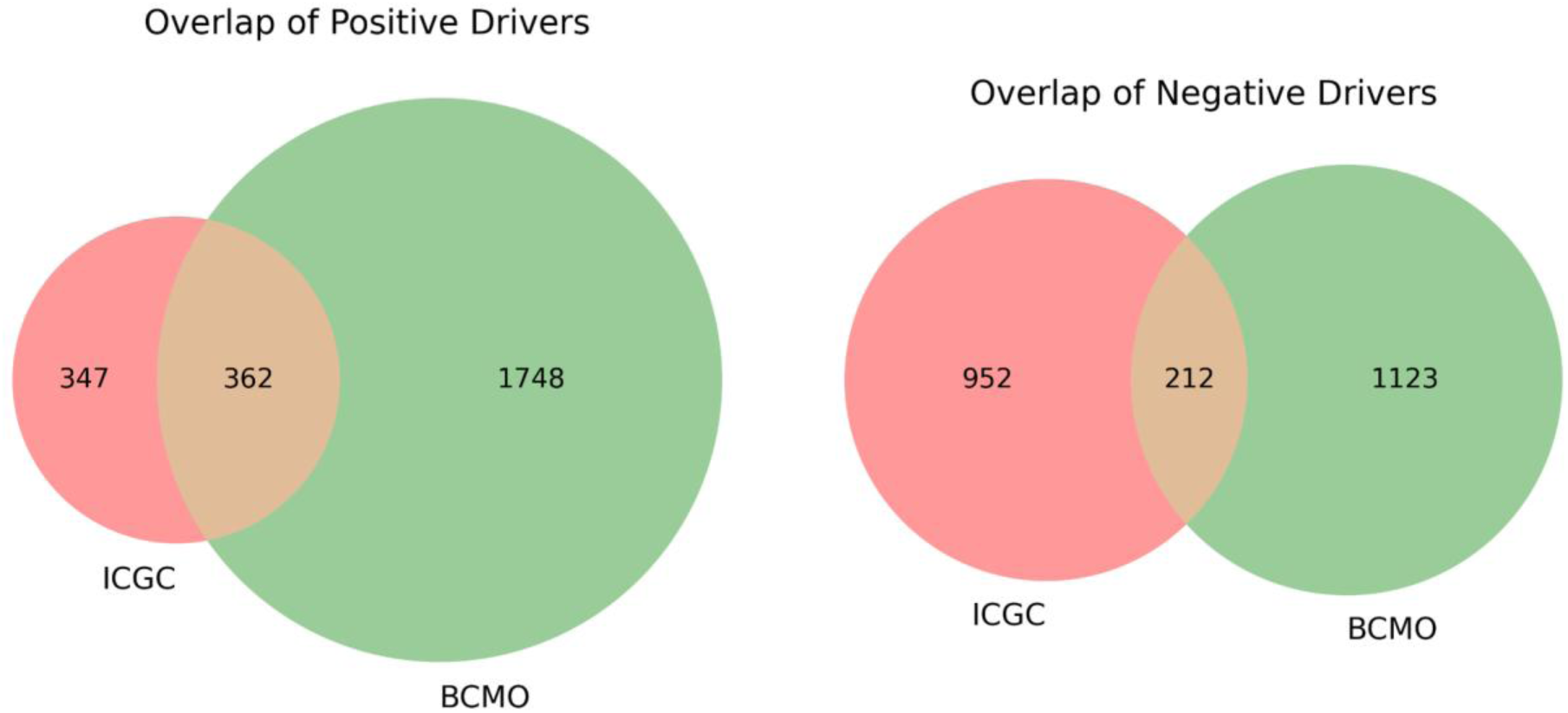
Venn plots indicating the overlapped positive and negative drivers across studies.

**Supplementary Figure 2.**
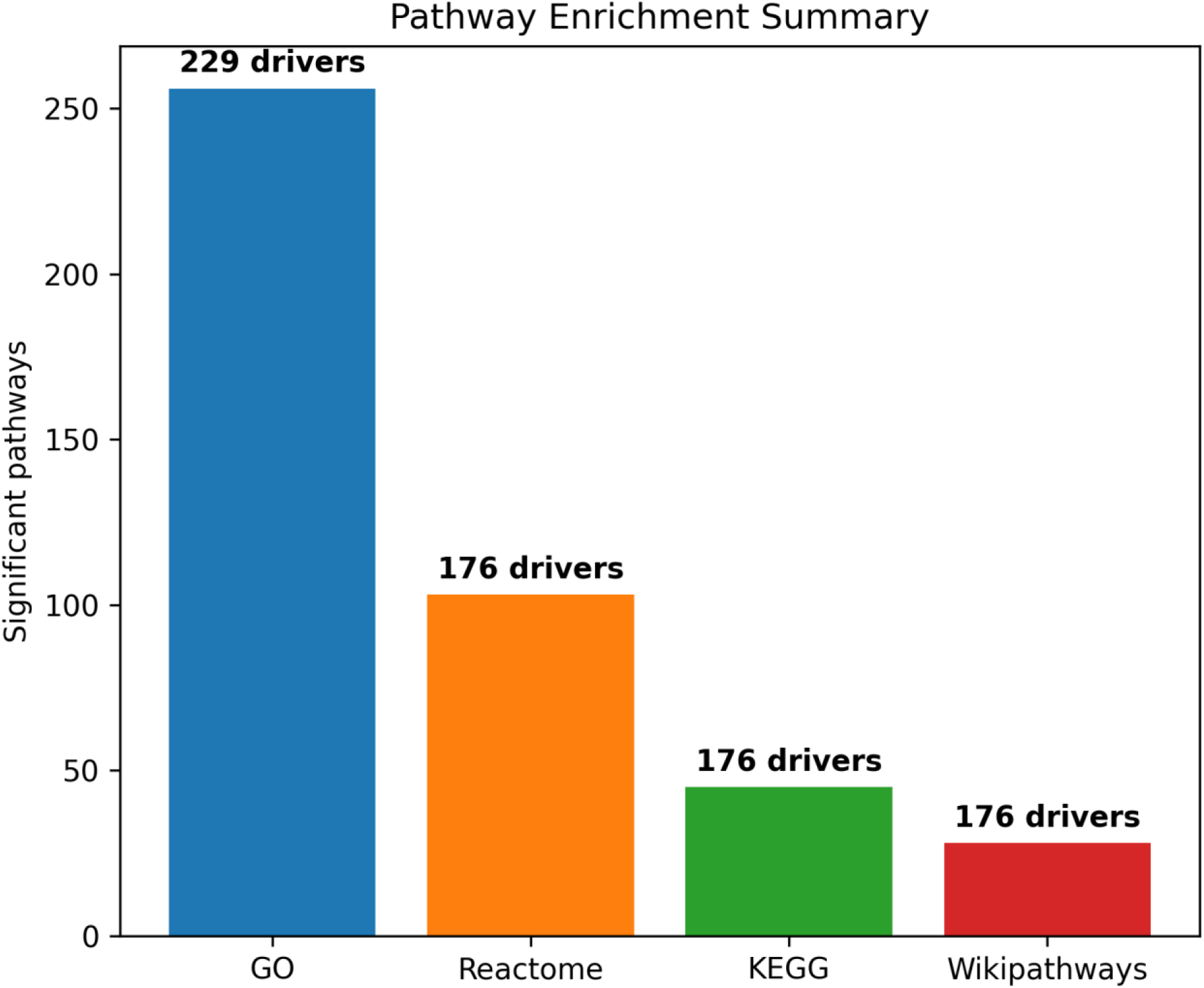
Number of significant pathways identified in each database through enrichment analysis for CLL case study. Bars represent the pathways, and the numbers above each bar indicate the number of associated drivers in each set.

**Supplementary Figure 3.**
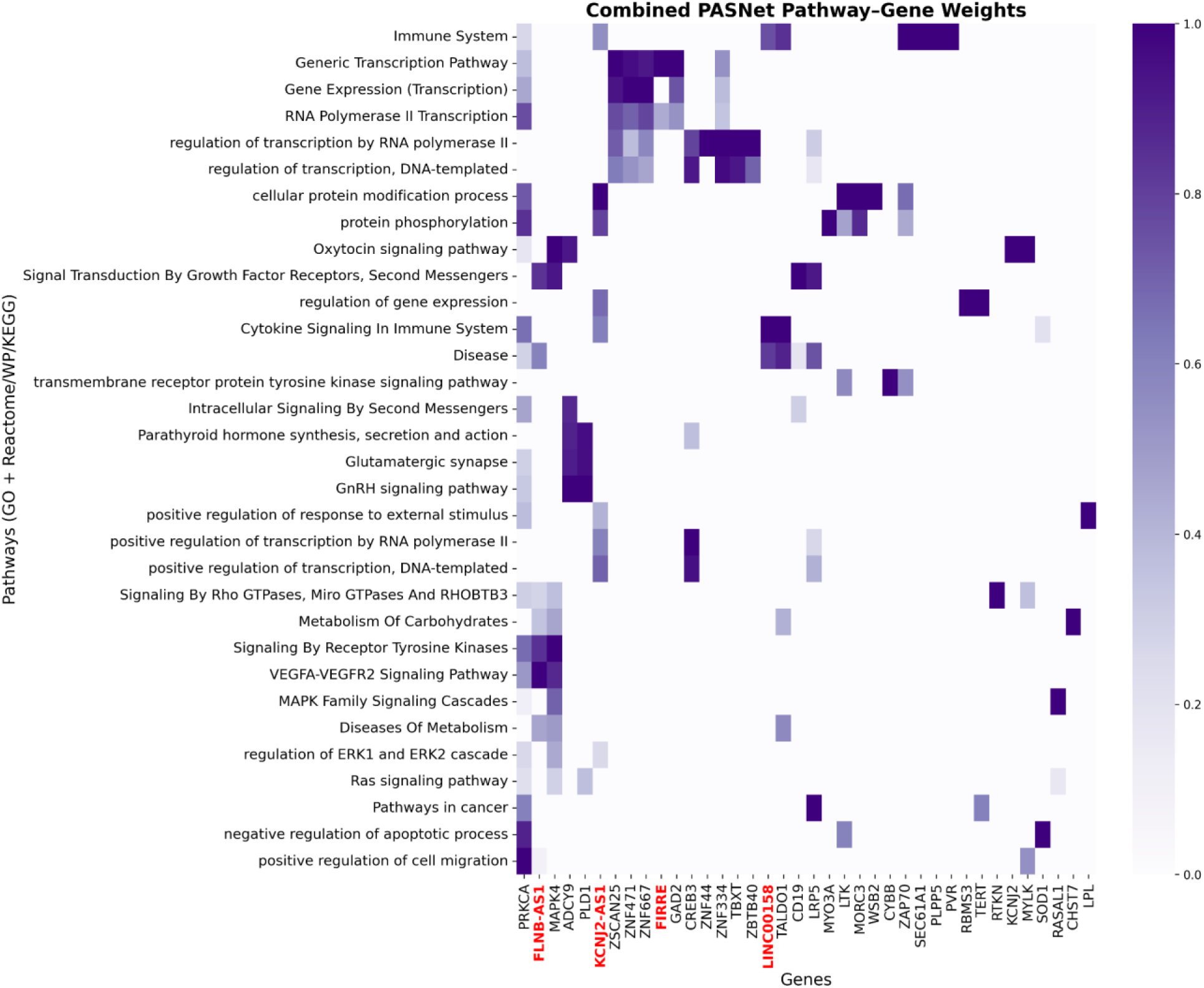
Heatmap depicting, combined from both models, weight correlations between pathways (connected to at least three driver genes) and top SHAP values from each model that connected to at least one pathway for the CLL case study.

**Supplementary Table 1.**
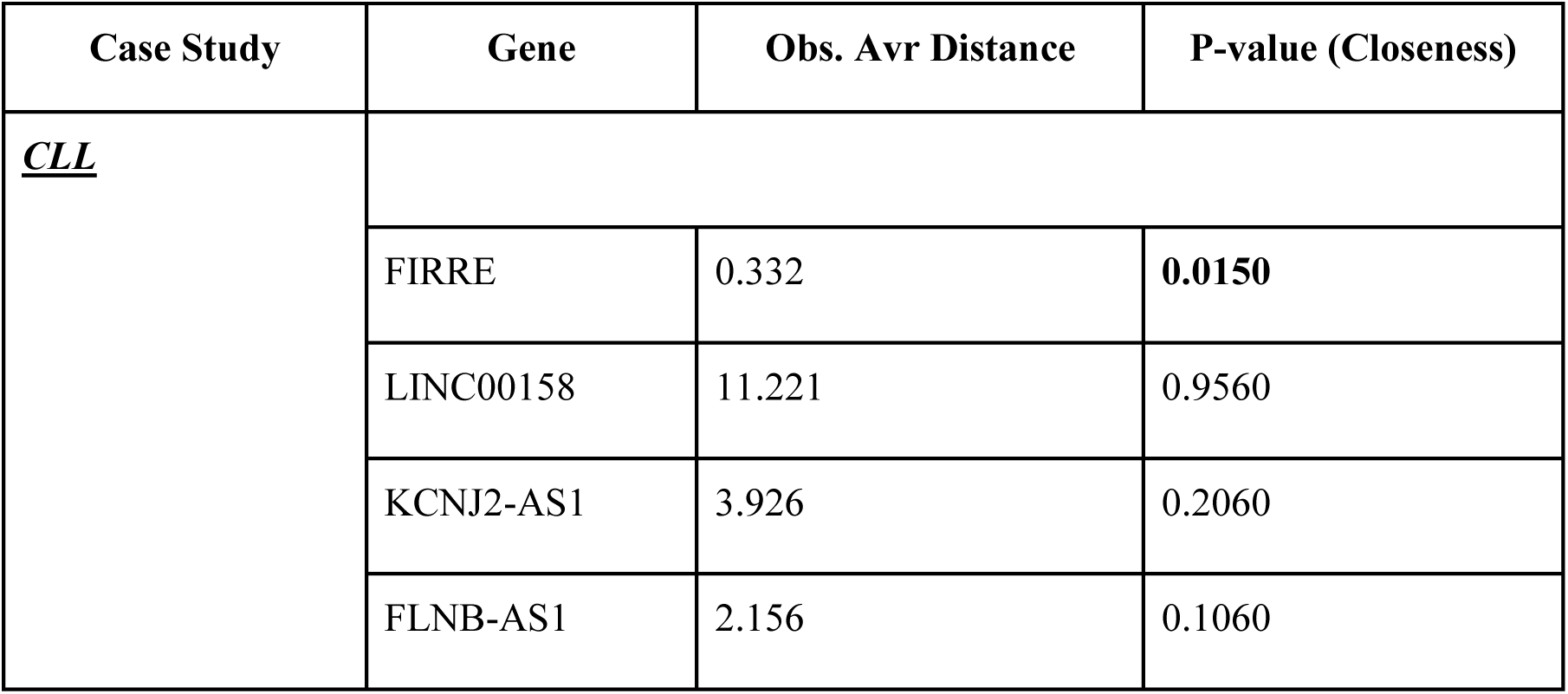
Summary of Gene Associations in CLL case study with Observed Average Distances Between lncRNAs Identified in Studies and Experimentally Validated lncRNAs from the lncHUB UMAP.

| Case Study | Gene | Obs. Avr Distance | P-value (Closeness) |
| --- | --- | --- | --- |
| <u>**CLL**</u> |  |  |  |
|  | FIRRE | 0.332 | **0.0150** |
|  | LINC00158 | 11.221 | 0.9560 |
|  | KCNJ2-AS1 | 3.926 | 0.2060 |
|  | FLNB-AS1 | 2.156 | 0.1060 |

**Supplementary Figure 4.**
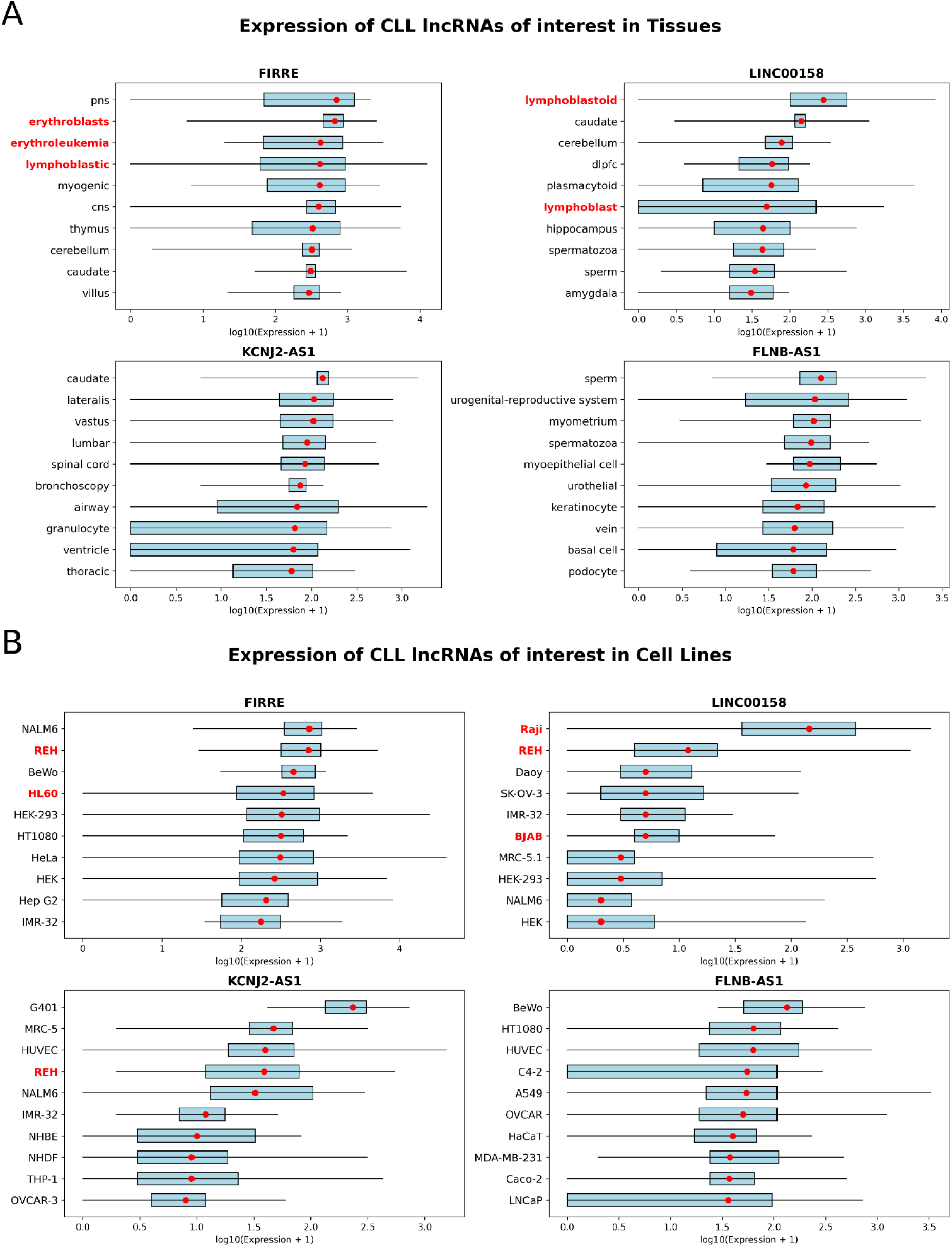
Box plots showing lncRNA expression across various tissues and cell types for the CLL case study scenario. For the CLL case study, the top 10 most highly expressed tissues (A) and cell types (B) for each lncRNA were extracted from ARCH4.

**Supplementary Figure 5.**
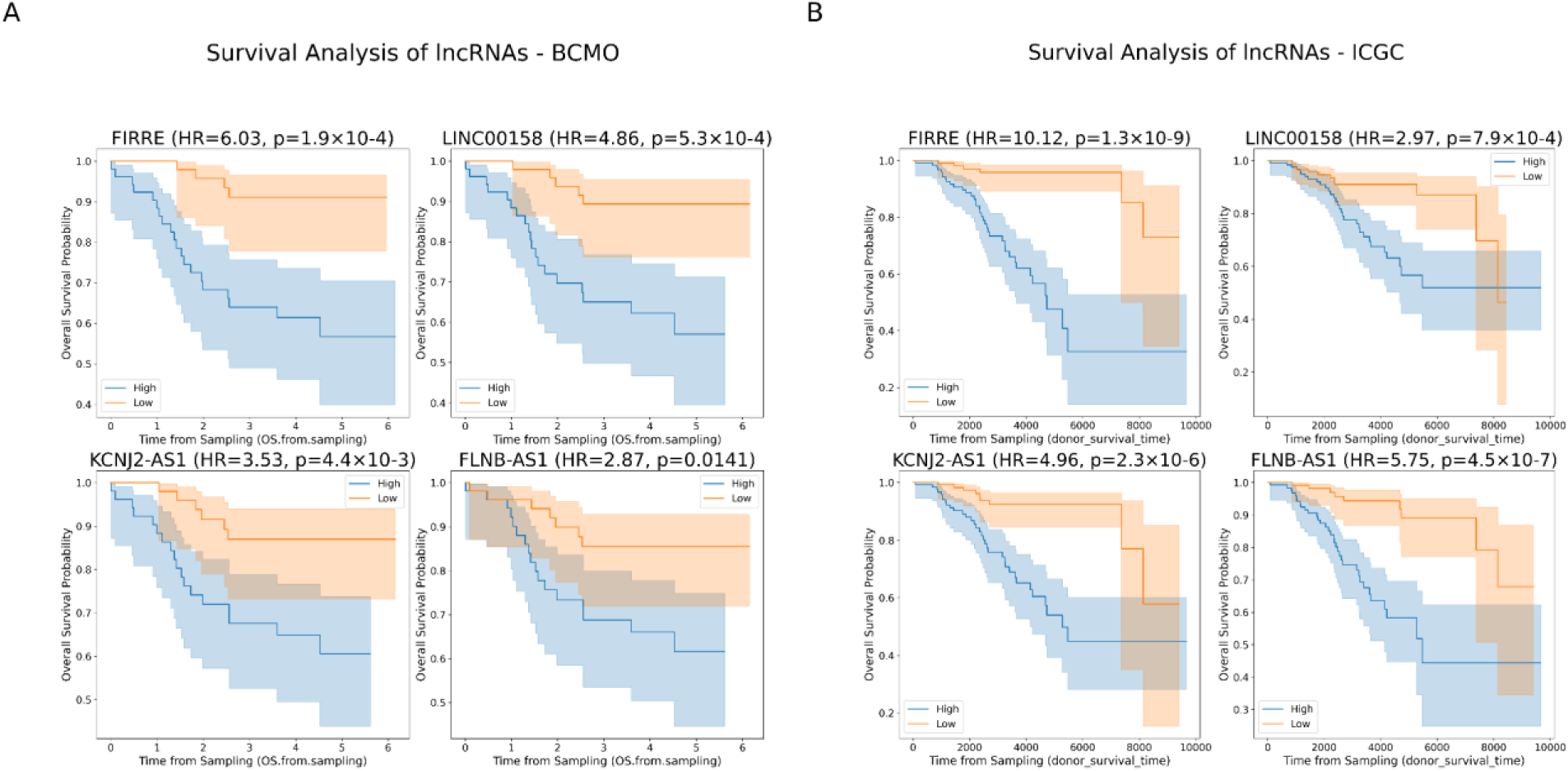
Kaplan–Meier plots for each significant lncRNA identified in the CLL analysis. Survival curves are shown for samples with high (blue) versus low (orange) lncRNA activity. Plots are presented for (A) the BCMO dataset and (B) the ICGC dataset.

**Supplementary Table 2.** Summary of Gene Associations in PRAD case study with Observed Average Distances Between lncRNAs Identified in Studies and Experimentally Validated lncRNAs from the lncHUB UMAP.

| Case Study | Gene | Obs. Avr Distance | P-value (Closeness) |
| --- | --- | --- | --- |
| **<u>Prostate</u>**<br>**<u>Adenocarcinoma</u>** |  |  |  |
|  | PRC1-AS1 | 0.200 | **0.0210** |
|  | FRY-AS1 | 0.751 | 0.2150 |
|  | LINC01082 | 0.284 | **0.0400** |
|  | HCG11 | 1.101 | 0.3830 |

**Supplementary Figure 6.**
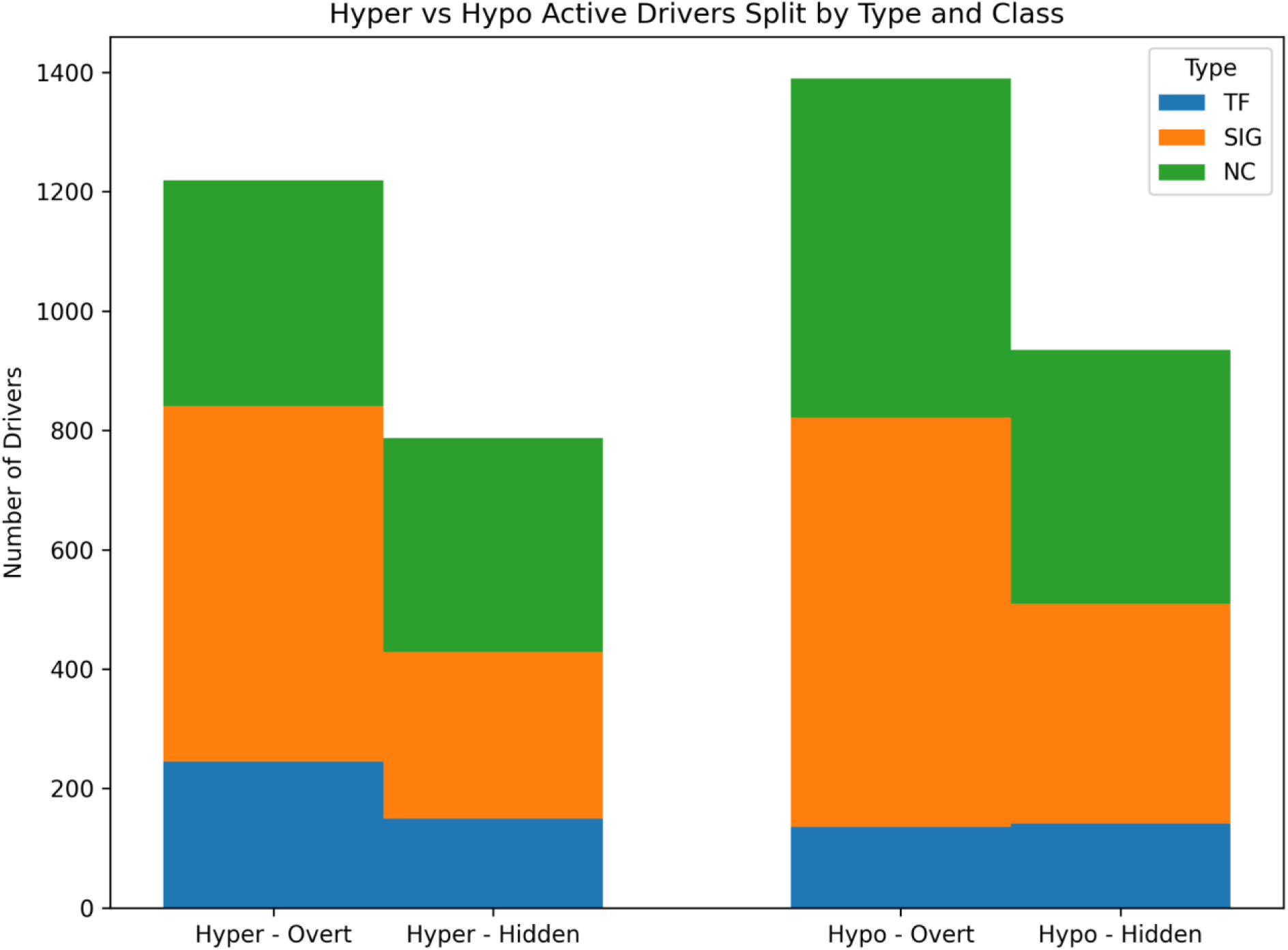
Distribution of Hyper-active and Hypo-active significant drivers in PRAD data. The drivers were separated by activity state (Hyper/Hypo), driver class (Overt vs Hidden), and type (TF (blue), SIG (orange), and NC (green)). Each bar represents the number of drivers in the corresponding category, with stacked colors indicating driver type.

**Supplementary Figure 7.**
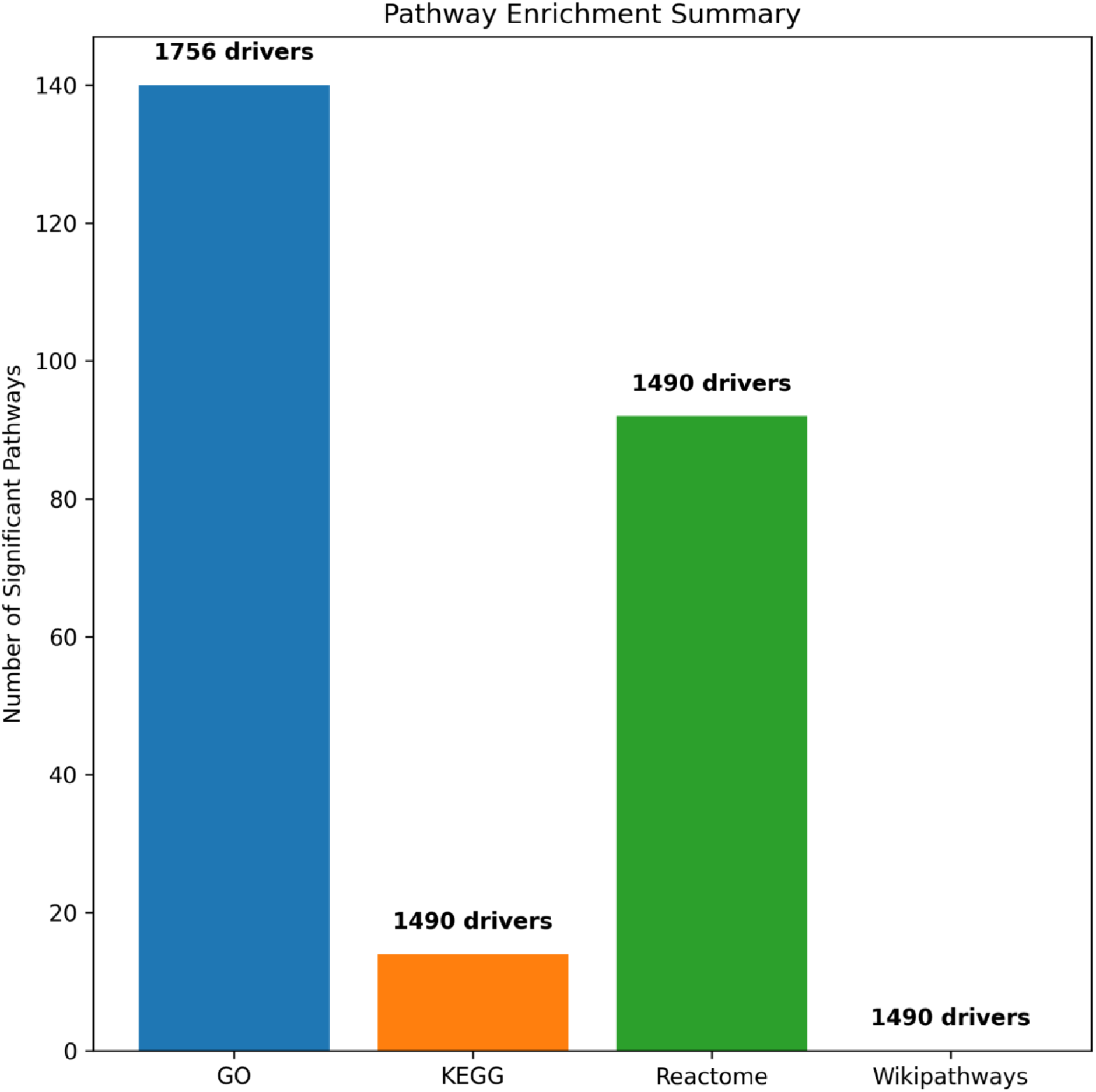
Number of significant pathways identified in each database through enrichment analysis for PRAD case study. Bars represent the pathways, and the numbers above each bar indicate the number of associated drivers in each set.

**Supplementary Figure 8.**
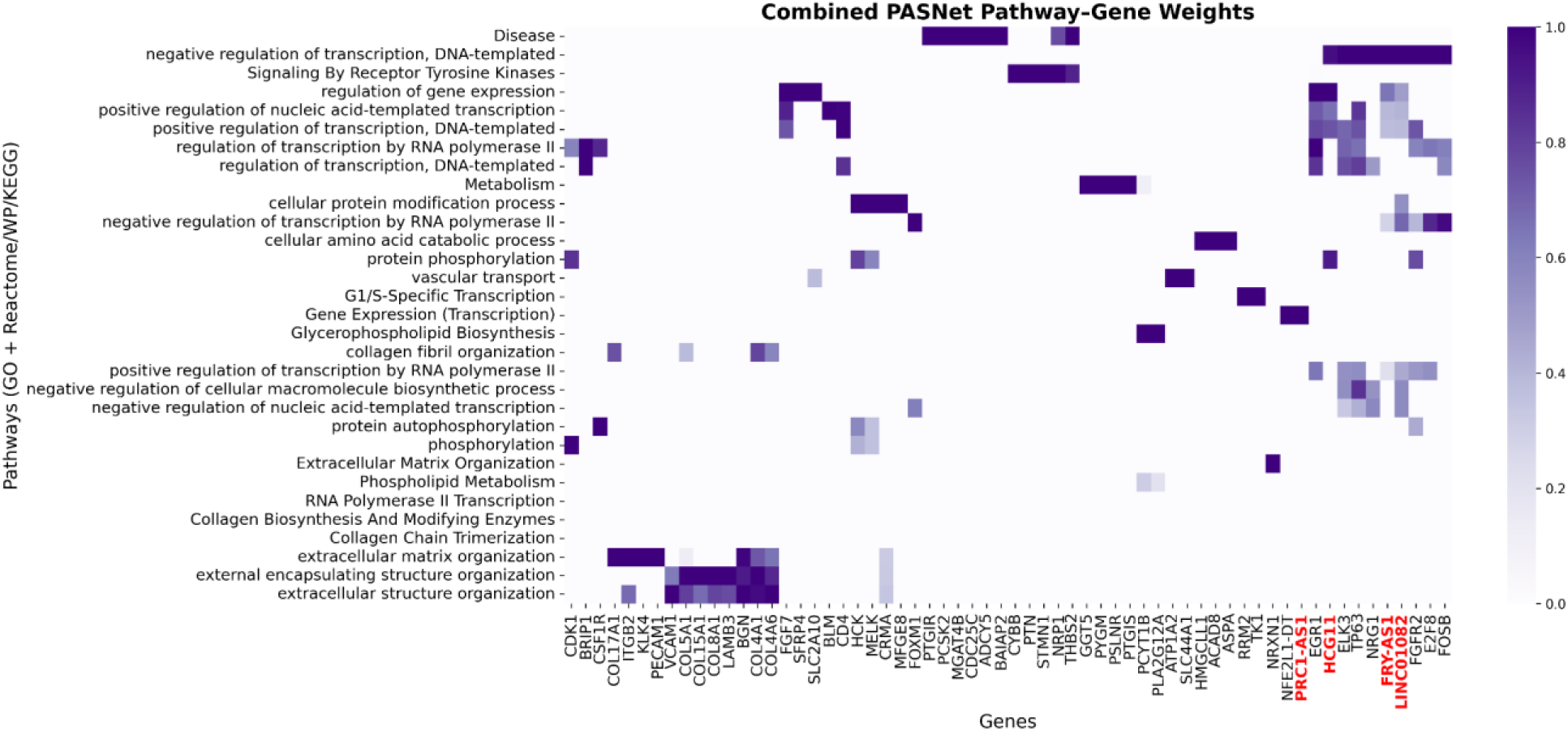
Heatmap depicting, combined from both models, weight correlations between pathways (connected to at least three driver genes) and top SHAP values from each model that connected to at least one pathway for the PRAD case study.

**Supplementary Figure 9.**
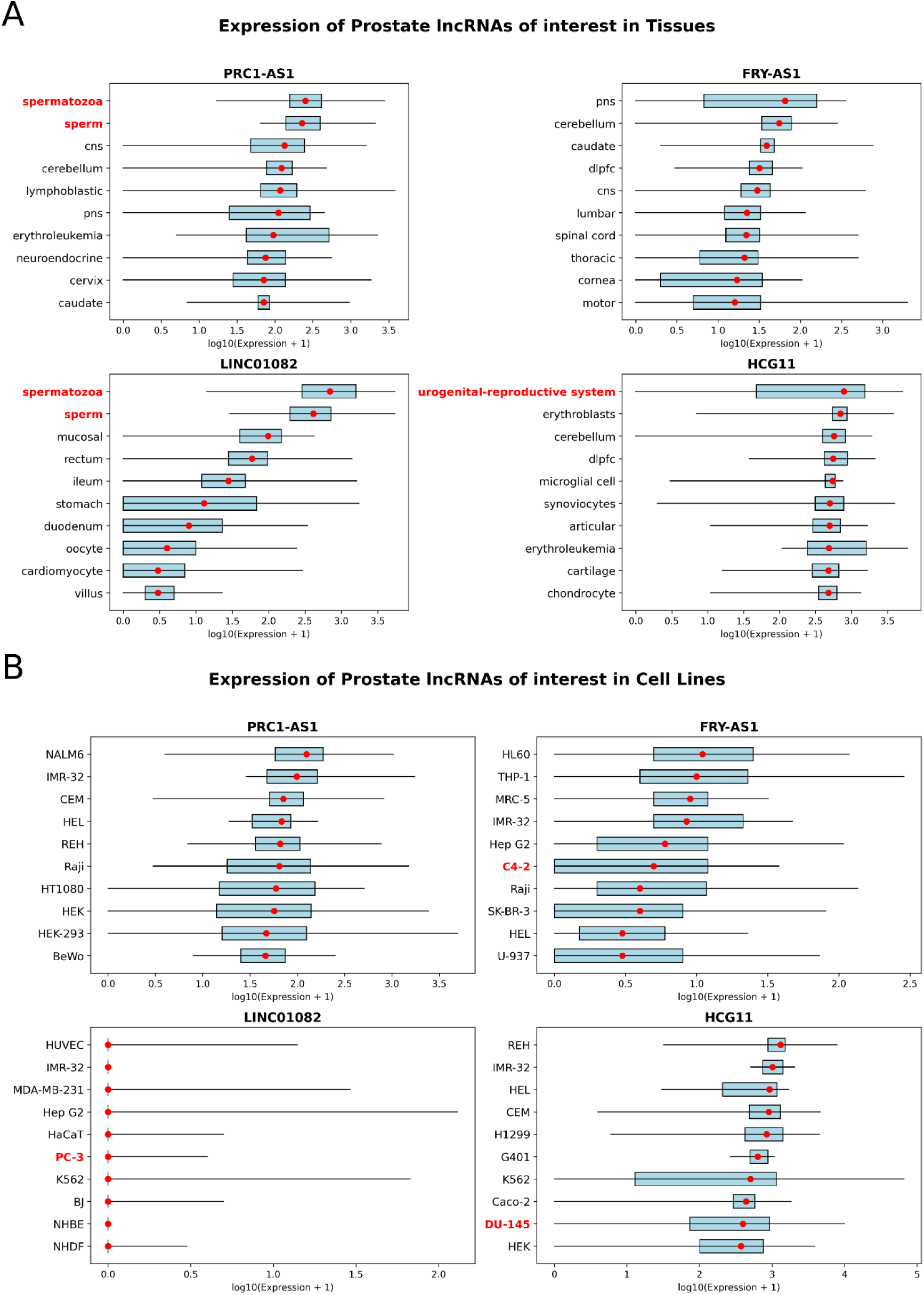
Box plots showing lncRNA expression across various tissues and cell types for the Prostate Adenocarcinoma case study scenario. For the PRAD case study, the top 10 most highly expressed tissues (A) and cell types (B) for each lncRNA were extracted from ARCH4.

